# Growing Minds, Integrating Senses: Neural and Computational Insights into Age-related Changes in Audio-Visual and Tactile-Visual Learning in Children

**DOI:** 10.1101/2025.03.04.639748

**Authors:** Nina Raduner, Carmen Providoli, Sarah V. Di Pietro, Maya Schneebeli, Iliana I. Karipidis, Ella Casimiro, Saurabh Bedi, Michael von Rhein, Nora M. Raschle, Christian C. Ruff, Silvia Brem

## Abstract

Multisensory processing and learning shape cognitive and language development, influencing how we perceive and interact with the world from an early age. While multisensory processes mature into adolescence, it remains poorly understood how age influences multisensory associative learning. This study investigated age-related effects on multisensory processing and learning during audio-visual and tactile-visual learning in 67 children (5.7–13 years) by integrating behavioural and neuroimaging data with computational methods. A reinforcement-learning drift diffusion model revealed that older children processed information faster and made more efficient decisions on multisensory associations. These age-related increases coincided with higher activity in brain regions associated with cognitive control, multisensory integration, and memory retrieval, specifically during audio-visual learning. Notably, the bilateral anterior insula exhibited heightened activation in response to lower reward prediction errors, indicative of increased sensitivity to negative feedback with development. Finally, reward prediction errors modulated activation in reward processing and cognitive control regions, with this modulation remaining modality-independent and largely stable across age. In conclusion, while children employ similar learning strategies, older children make decisions more efficiently and engage neural resources more strongly. Our findings reflect ongoing maturation of neural networks supporting multisensory learning in middle childhood, enabling more adaptive learning in later childhood.

**Highlights:**

– faster information processing in older children during a multisensory learning task
– increasing brain activation with age in visual, parietal, and frontal regions
– reward prediction error processing are independent of age and modality
– heightened response to negative reward prediction errors with increasing age

## 1 Introduction

Efficient processing and integration of sensory information from multiple modalities including vision, hearing, and touch, are essential for interacting with our surroundings and adapting to the complexities of our environment. On a daily basis, humans encounter numerous stimuli by various senses, requiring appropriate processing and integration to be accurately understood (Van Atteveldt et al., 2014). Multisensory processing describes the perception of stimuli in multisensory environments (Murray et al., 2016). Multisensory learning (MSL) is the process of acquiring associations between stimuli from different senses (Lauzon et al., 2022), while multisensory integration (MSI) occurs when the brain combines information from multiple senses to form a unified percept (Stein et al., 2020; Wallace et al., 2020). When stimuli from different modalities provide congruent information, they are integrated and perceived as originating from the same source. Research using electrophysiology, fMRI, and behavioural studies has shown that successful associative MSL leads to MSI (Butler et al., 2011; Lauzon et al., 2022). Since these processes are sequential (multisensory processing is needed for MSL which is also the basis for MSI), they are often studied in one framework. MSI is particularly significant in naturalistic learning environments and classrooms, where multisensory exposure is common (Barutchu et al., 2019). Moreover, MSI skills are associated with enhanced cognitive processes including memory performance and reading abilities (Barutchu et al., 2011, 2019; Birch & Belmont, 1965; Denervaud et al., 2020; Dionne-Dostie et al., 2015). MSI likely promotes learning by enabling the effective combination of sensory inputs for better encoding, storage, and retrieval of information. For example, MSI underpins language learning, production, and development by integrating auditory and visual information like sounds and lip movements (Wallace et al., 2020).

Given the importance of MSL and MSI in cognitive development, it is crucial to investigate MSL in childhood and how it is similar or different across different sensory modalities. However, most studies on MSL have focused on adults and the studies investigating MSL in children mainly concentrated on audio-visual speech sound mapping (e.g., Frei et al., 2025; Karipidis et al., 2021; Wang et al., 2020). Only a few investigations targeted the learning and processing of non-linguistic audio-visual integration (e.g., Altarelli et al., 2020; Batson et al., 2011; Pasqualotto et al., 2024; Seitz et al., 2007). Moreover, while MSL can encompass signals from all senses, minimal attention has been directed towards the interplay between touch and vision or any other combination of stimulus modalities (see meta-analysis from Gao et al., 2023).

Despite the growing interest in and the importance of MSL, relatively few studies have examined its neural underpinnings (e.g., Butler et al., 2011; Lauzon et al., 2022), with most research focusing on MSI or failing to distinguish between the two processes. The meta-analysis by Gao et al. (2023) further highlights that audio-visual integration has been widely studied, whereas only a few studies examined tactile-visual integration (such as Gentile et al., (2013, 2015); Limanowski & Blankenburg, (2016, 2017)). In the mentioned meta-analysis, audio-visual integration was associated with changes in neural activation of early sensory regions, subcortical areas, and higher association areas supporting a flexible neural pathway model for audio-visual integration. Such a model is supported by research linking MSI and learning to an extended cortico-striato-thalamic network (Lakatos et al., 2007; Murray et al., 2016; Tyll et al., 2011), including early sensory regions (Ghazanfar & Schroeder, 2006; Lakatos et al., 2007; Noesselt et al., 2007; Stein & Stanford, 2008). Moreover, reviews summarising brain regions involved in MSI identified similar regions to be relevant for these processes: the auditory cortex, visual cortex, somatosensory cortex, superior temporal cortex, and frontal regions (Murray et al., 2016; Richlan, 2019). Exploratory meta-analyses on the few existing studies on tactile-visual integration reveal that inferior temporal areas, the fusiform gyrus, and inferior parietal areas are associated with tactile-visual integration (Gao et al., 2023). This highlights the need for further research on the brain areas underlying MSL, particularly during childhood, when these abilities are still developing.

Although infants already demonstrate the ability to process and integrate multisensory information (Patterson & Werker, 1999, 2003), a deeper understanding of how these abilities develop over childhood is crucial for advancing learning research. MSI has not yet reached full maturity in children younger than eight years old and maturation of MSI skill continues into adolescence (Desjardins et al., 1997; Dionne-Dostie et al., 2015; Gori et al., 2008; Nardini et al., 2008). Research suggests the temporal window for audio-visual integration is still immature in 11-year-old children (Dionne-Dostie et al., 2015). According to Murray et al. (2016), children’s MSI abilities are initially more reliant on low-level stimulus feature processing but gradually shift toward higher stimulus feature processing and therefore cortical involvement. This process reflects the brain’s developmental trajectory where subcortical regions mature before cortical regions. This implies that younger children, who depend more on the processing of low-level features through subcortical and primary sensory areas, might have limited MSI capabilities for complex learning tasks requiring greater levels of cognitive control. With children’s increasing age, however, the integration of sensory information progressively involves higher-order brain regions, such as the frontal cortex. This in turn allows for more flexible and sophisticated MSI that is crucial for advanced learning, problem-solving, and adaptation in dynamic environments. As a result, these maturation processes allow for more complex multisensory learning and integration as children grow (Murray et al., 2016).

To better understand the developmental processes, computational modelling provides a powerful tool for examining the cognitive mechanisms underlying MSL, deepening the neural and developmental insights. One approach in computational modelling integrates a learning process, reflecting trial-wise belief updating, and a choice process, translating computational quantities (e.g., values) into behavioural or neural responses (Mathys et al., 2011). This study employs a combined reinforcement learning model as the learning process and drift diffusion model as the choice process (RLDDM; Pedersen et al., 2017) to investigate multisensory processing and learning in children (see also Fraga-González et al., 2025; Frei et al., 2025). Reinforcement learning, one of the core mechanisms for feedback learning, compares observed outcomes (rewards) with expected outcomes (values) and adjusts expectations based on the discrepancy – known as the reward prediction error (RPE; Niv & Schoenbaum, 2008; Subramanian et al., 2022). The Rescorla-Wagner (RW) model describes this process by weighting the RPE with the learning rate to update values (Rescorla & Wagner, 1972). On the other hand, the drift diffusion model quantifies the decision-making process through the drift rate, decision boundary, and non-decision time (Pedersen et al., 2017; Ratcliff, 1978; Ratcliff & Rouder, 1998). By combining these models, we can identify latent cognitive processes like value updating, learning rates, and RPEs, and relate them to the brain activation during learning.

Building on these computational frameworks, previous research has identified specific neural correlates of the cognitive parameters involved in learning. RPEs are consistently linked to stronger brain responses in the striatum, the medial frontal cortex, and hippocampus reflecting their relation to learning and memory (Cohen & Ranganath, 2005; Fraga-González et al., 2025; Garrison et al., 2013; McClure et al., 2003; O’Doherty et al., 2003; Pagnoni et al., 2002). Similarly, value processing involves regions such as the ventral striatum, anterior cingulate cortex, anterior insula, medial orbitofrontal cortex, supplementary motor area, and inferior parietal lobe (Liu et al., 2011; R. P. Wilson et al., 2018). These regions are pivotal in integrating motivational and emotional aspects of decision-making (Liu et al., 2011). The learning rate has been linked to activation in the inferior frontal gyrus, anterior insula, dorsal anterior cingulate cortex, and ventral caudate (Chien et al., 2016; O’Reilly, 2013; Wu et al., 2017). By mapping neural correlates of computational parameters such as values, reward prediction errors, and learning rates, our study aims to computationally characterize the perceptual and learning processes underlying MSL in children and study their change during development.

To further explore these neural and cognitive processes, the current study investigated MSL in middle childhood, focusing on learning sensory associations of different modalities (audio-visual vs. tactile-visual) and on age-related changes therein. The primary aims were: 1) to **investigate modality-specific differences in children’s multisensory processing and learning** by comparing audio-visual (AV) and tactile-visual (TV) associations, using computational modelling of behavioural performance and neural activity data, 2) to characterize the **age-related changes in MSL** by analysing age-related differences in processing speed, decision-making and associated neural activity, and 3) **to examine how children process feedback during MSL**, specifically looking at age-effects on RPE and value-based learning for audio-visual and tactile-visual learning. We hypothesised that younger children would rely more on subcortical and primary sensory regions, while older children would engage higher-order cortical areas during MSL, indicating the increased employment of neural resources (Murray et al., 2016). Accordingly, we anticipated neural activation in brain regions previously associated with value processing (Liu et al., 2011; R. P. Wilson et al., 2018), reward prediction error processing (Cohen & Ranganath, 2007; Fraga-González et al., 2025; Garrison et al., 2013; McClure et al., 2003; Pagnoni et al., 2002), and learning rate processing (Chien et al., 2016; O’Reilly, 2013; Wu et al., 2017). We expected potential age-dependent variations likely to be pronounced in higher-cortical areas such as prefrontal and cingulate regions (Yaple et al., 2020). With this study, we bridge the gap in the existing literature by investigating non-linguistic audio-visual and tactile-visual associative learning across middle childhood.

We employed a cross-sectional design with 67 children (aged 5.7–13.0 years) completing a multisensory learning task involving AV and TV stimuli. Computational modelling, incorporating reinforcement learning and drift diffusion frameworks, was applied to characterise the cognitive processes underpinning learning. Functional MRI was employed to pinpoint brain regions engaged in modality-general and modality-specific learning processes and to characterize age-related changes in the activation of involved brain networks. Parametric modulation of the BOLD signal with model parameters finally allowed us to examine the neural correlates underlying specific learning and decision processes, as well as potential age-related changes in the neural processing.

## 2 Material and methods

### 2.1 Participants

This study included 80 healthy, native (Swiss-)German-speaking children. Children with either neurological or psychiatric disorders or intellectual disabilities (except for specific learning disorders or ADHD) were not included to participate in this study. Following the exclusion of one subject due to technical problems during scanning, another two subjects had to be excluded due to excessive movement in all runs (see chapter 2.3) and a high number of omissions (see Supplementary Material A.6). Additionally, 10 children were excluded from the final analysis as they had only one AV or TV run with sufficient MR data quality (no excessive movement). Consequently, the final sample comprised 67 children (*M* = 8.35 ± 1.82 years, 36 girls) with at least one high-quality run in both the AV and TV modalities. There was no significant age difference between girls and boys (see Supplementary Material A.1). All children had estimated IQ scores above 80. All participants had normal or corrected to normal vision and normal hearing. Mean IQ, processing speed, and attention scores were in the normal range (see **Table 1**; a table summarizing the demographics of the group split in 3 age bins [5.7-8.0, 8.1-10.5; 10.6-13] can be found in the Supplementary Material A.1), indicating age-appropriate cognitive abilities for all subjects (see Supplementary Material A.1 for more information). A total of ten children had either a clinical diagnosis (attention deficit hyperactivity disorder, developmental dyslexia, or developmental language delay) and/or clinically significant values in the CBCL questionnaire (Döpfner et al., 2014; see Supplementary Material A.1 for more information). The following analyses were conducted with and without these ten participants (see Supplementary Material B.1). This approach ensures that the results are not biased by potential neurodevelopmental differences while also allowing for a more comprehensive analysis that includes all participants.

**Table 1.** Demographic characteristics of study participants.

|  | *M (SD) [range]* |
| --- | --- |
| **Age [y]** | 8.35 (1.82) [5.7-13] |
| **Sex [f:m]** | 36:31 |
| **Handedness<sup>a</sup> [l:a:r]** | 6:3:57 |
| **IQ (verbal and non-verbal)** | 106.5 (10.6) |
| **Rapid automatized naming [T-value]** | 50.1 (7.45) |
| **Processing speed [T-value]** | 50.12 (7.97) |
| **Sustained attention median RT<sup>b</sup> [PR]** | 51.55 (29.43) |

*Note.* The handedness of the subjects was determined using the Edinburgh Handedness Scale (Oldfield, 1971). IQ values were estimated using the Peabody Picture Vocabulary Test – fourth edition (PPVT-IV; Lenhard et al., 2015) and subtests from the Wechsler Intelligence Scale for Children – fifth edition (WISC-V; Wechsler, 2017) or the Wechsler Preschool and Primary Scale of Intelligence – fourth edition (WPPSI-IV; Wechsler, 2012). Rapid automatized naming and processing speed measured with IDS-2 (Grob & Hagmann-von Arx, 2018). Sustained attention measured with KiTAP (Zimmermann et al., 2002). <sup>a</sup> One child's data are missing. <sup>b</sup> Percentiles are missing for two children due to the absence of normative values for their age. y = years, f = female, m = male, l = left, a = ambidextrous, r = right, RT = reaction time, PR = percentiles.

Oral consent was obtained from all children, while written consent was provided by a parent or guardian for all children. The study was approved by the local ethics committee of the Canton of Zurich and neighbouring cantons in Switzerland (BASEC-Nr: 2022-01368), and the participants received vouchers and gifts in appreciation of their participation.

### 2.2 Experimental design

During the neuroimaging session, participants completed an audio-visual and tactile-visual multisensory learning task (adapted from Frei et al. (2025)) embedded in a child-friendly story in the MRI scanner. The multisensory learning task is a two-alternative forced choice task where the participants had to learn multisensory associations between visual and auditory or visual and tactile stimuli based on feedback. Participants were presented with two symbols alongside either a sound or vibration pattern, representing a rocket’s sound or engine trembling, respectively. Participants guessed which rocket the sound or trembling belonged to by pressing a corresponding button. They were told to learn associations over time through feedback, which was implicitly suggested to be probabilistic, and to collect as many stars as possible (rewarded after a correct response; incentive feedback) during each run. The task was presented using goggles (CinemaVision 20/20, Resonance Technology, Northridge, CA), MR-compatible over-ear headphones (MR confon GmbH, Magdeburg, Germany, http://www.mr-confon.de/de/), a tactile stimulation device (mini-PTS piezoelectric stimulator device from dancer design, http://www.dancerdesign.co.uk/) stimulating the right pinkie, and the PsychoPy® software (Version 2022.1.1, Peirce et al. (2019); https://www.psychopy.org/). To ensure that each child could perceive the stimuli clearly during the MR session, the sound volume was adjusted, and vision was corrected individually as needed.

Not all children were able to complete every task according to the experimental protocol and therefore runs had to be repeated or excluded due to excessive movement, lack of motivation (too many omissions, see Supplementary Material A.6), or fatigue. For the purposes of this analysis, only data from children who completed at least one audio-visual and one tactile-visual run with adequate data quality were included. The 67 participants included in these analyses finally showed the following distribution across the task modalities: 18 children completed two runs of each modality, 39 completed only one run of each, 5 completed two AV and one TV run, and 5 completed one AV and two TV runs.

As depicted in **Figure 1**, during the inter-trial interval (ITI), the task background consisted of a space image (dark background with stars), two blank rockets on either side of a larger star in the middle, acting as the fixation cross. These elements were constantly displayed during the whole task. During stimulus presentation, one auditory stimulus was presented simultaneously with two visual stimuli appearing on either rocket for a duration of 2 seconds (**Figure 1**). In TV runs, a tactile stimulation was presented instead of an auditory sound (see Supplementary Material A.2). The participants were then instructed to choose the matching symbol by pressing the left or right button on an MR-compatible two-button response box for the left or right choice, respectively, and their choice was indicated by a blue frame during the inter-stimulus interval (ISI). The participant’s choice was recorded during stimulus presentation and the ISI. After the response, they received feedback on their choice for 1.8s, which was followed by an ITI before the next trial began. The feedback in the task was probabilistic, with correct (e.g., positive feedback after a correct choice) feedback in most trials, and some trials providing incorrect (e.g., negative feedback after a correct choice; see Supplementary Material A.3 for more information). Neutral feedback was given after no choice was observed (see Supplementary Figure A.2B). The ITI and ISI were gamma distributed and had a mean duration of 1.87s [0.76 – 3.45s] and 3.45s [2.28 – 4.83s], respectively. After the first six participants, we slightly changed the timing of the ISI and ITI to provide more decision time. For these participants, the mean durations were 2.06s [0.79 – 3.39s] and 2.67s [1.11 – 4.34s], respectively for the ITI and ISI. Each run consisted of 44 trials and total task duration was 401 seconds (384 seconds for the first 6 subjects).

**Figure 1.**
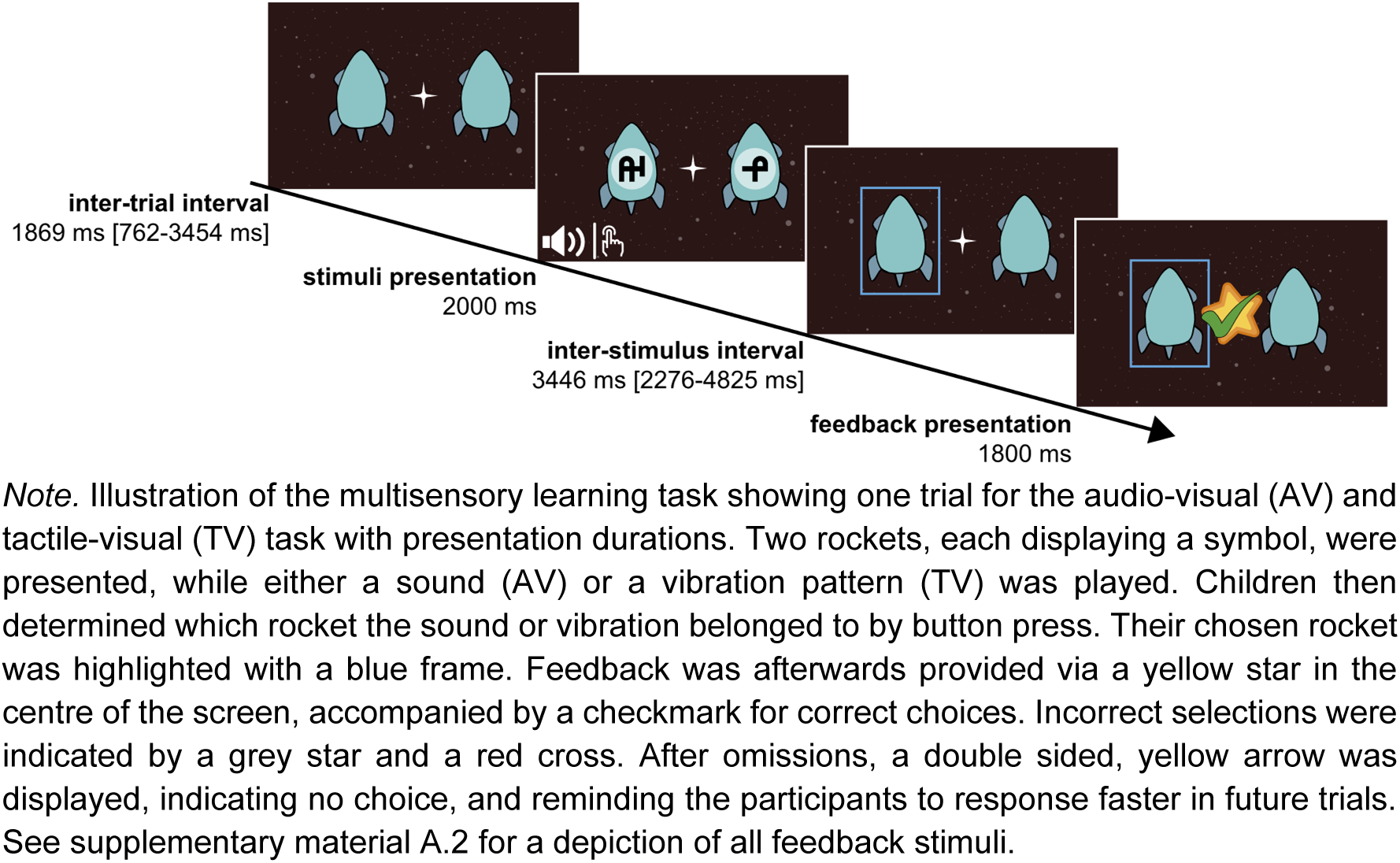
Multisensory learning task.

In total, there were four visual stimuli and four auditory or tactile stimuli per run, resulting in 16 possible combinations of which four were matching combinations. For the MR runs, we prepared six sets of visual, three sets of auditory stimuli, and three sets of tactile stimuli (see Supplementary Material A.2). As visual stimuli, artificial symbols from the PseudoSloan font were used (Vildavski et al., 2021). The pseudo letters were chosen by the complexity scores provided by Vildavski et al. (2021). Each set was matched such that it included one stimulus with low (*M* = 3.55, *SD* = 0.02), two stimuli with medium (*M*_1_ = 4.27, *SD*_1_ = 0.09 and *M*_2_ = 4.49, *SD*_2_ = 0.05) and one stimulus with high complexity (*M* = 5.16, *SD* = 0.05). Additionally, we also balanced the selection of pseudo letters between sets. For the audio-visual run, environmental sounds were used as auditory stimuli, which were selected from the mixkit sound effects database (https://mixkit.co/free-sound-effects/). Here, we chose sounds that could not be easily identified or named. After selection, the sounds were cut to a 2 second duration and normalized using Audacity® recording and editing software (version 3.5.1; Team Audacity, 2024). The mean Shannon spectral entropy (Cover & Thomas, 2005; Misra et al., 2004; Obin & Liuni, 2012), a measure of sound complexity, was calculated for each stimulus using the spectralEntropy function (Signal Processing Toolbox, version 9.1) in MATLAB (version R2022b; MathWorks Inc, 2022). Each set of environmental sounds consisted of one sound with a low (*M* = .327, *SD* = 0.03), two with a medium (*M*_1_ = .476, *SD*_1_ = 0.05 and *M*_2_ = .564, *SD*_2_ = 0.04), and one with a high entropy value (*M* = .705, *SD* = 0.04). Lastly, we created three different sets for the tactile stimulation. Each set of tactile stimuli consisted of four different vibration patterns lasting for 2 seconds. Each set contained two stimulations with distinct stable frequencies (low complexity), one was increasing or decreasing (medium complexity), and one had an (inverted) u-shape (high complexity; see Supplementary Figure A.2C). Ensuring that the stimuli varied based on complexity within a set was done to increase the discriminability between stimuli. The following variables were pseudo-randomised for all participants: the order of tasks (AV and TV) in the MR scanner, the trial structures (the sequence of stimulus presentation), the visual, auditory, and tactile sets used, and the assignment of the stimuli to the four correct combinations for each run.

Each participant completed behavioural runs (one for each modality) in a behavioural assessment session 1 – 42 days (*M* = 10.6 days, *SD* = 8.1 days) before the MR session (pre runs). During the behavioural session, each participant was walked through 13 instruction trials for the AV run by the examiner, afterwards they completed 10-20 practice trials on their own until the participant understood how the task works. Then they completed the first full AV behavioural run with 44 trials. Afterwards, the 13 instruction trials were repeated for the TV runs to familiarise the participants with the tactile vibration. If necessary, the 10-20 practice trials were also repeated, until the participants understood the task. Finally, each participant competed a full TV behavioural run with 44 trials. This procedure ensured that the participants were familiar with the task design before the MR session. The performance from the pre runs was then used to adapt the task difficulty in the MR scanner to ensure that the task was neither too easy nor too difficult (see Supplementary Material A.3 for adaptation criterions). Before entering the MR scanner, the task instructions were repeated to each participant, and they completed another 10-20 practice trials with the response box used in the MR. Therefore, each participant completed at least 134 trials of the multisensory learning task before entering into the MR. Importantly the stimulus material used for the introduction, practice, and pre-runs was different from the material used in the fMRI runs. Task difficulty was either adapted by the trial structure or by the level of probabilistic feedback. Additionally, some of the stimuli triplet combinations were presented more often than others (see Supplementary Material A.3 for more information about trial structure, probabilistic feedback, and stimulation frequency).

### 2.3 MRI data acquisition and preprocessing

MRI data was acquired on a Philips Achieva 3 Tesla scanner (Best, The Netherlands) using a 32-channel coil. T2* weighted functional images were acquired with a whole-brain gradient echoplanar image (EPI) sequence (five dummy scans followed by 291 dynamic scans, repetition time TR = 1.395 s, echo time TE = 35 ms, 44 slices, voxel size = 3.0 x 3.0 x 3.0 mm3, slice gap = 0.3 mm, matrix size 64 x 62 px, flip angle = 80°, multiband factor = 2, sofTone factor = 2, sensitivity-encoding (SENSE) reduction factor = 2). Due to changes in the timing of the task (see chapter 2.2), the EPI sequence of 6 participants only included 278 dynamic scans while all the other parameters were the same. T1-weighted anatomical images were obtained for co-registration using a magnetization-prepared rapid acquisition gradient echo (MPRAGE) sequence (TR = 6.96 ms, TE = 3.2 ms, aligned at AC-PC, flip angle = 9°, voxel size = 1.0 x 1.0 x 1.0 mm^3^, field of view = 270×255 mm^2^, number of slices = 176).

MATLAB (version R2020b, MathWorks Inc, 2020) toolbox SPM12 (7219, Penny et al., 2007) was used for preprocessing and whole-brain analysis, which included slice-time correction, realignment, co-registration, and segmentation. Using the Template-O-Matic toolbox (Wilke et al., 2008), a paediatric template was created for normalization (age matched structural data). Voxels were resampled to 3 x 3 x 3 mm^3^ isotropic voxels. An 8-mm full width at half maximum (FWHM) Gaussian kernel was applied for smoothing. After preprocessing, we repaired volumes with a composite scan-to-scan motion above 1.5mm (Euclidean distance with a radius of 65 mm) with the ArtRepair toolbox (Mazaika et al., 2007), using linear interpolation between the nearest unrepaired scans. After this, we flagged volumes with and without movement surrounded by scans with motion above 1.5mm. Runs with more than 20% of flagged volumes were excluded from the final analyses, resulting in a total of 18 runs being omitted (movement was not correlated with age; see Supplementary Material B.9). Whenever possible, we attempted to repeat runs with excessive movement during the MR session. Flagged scans were accounted for by including one binary regressor of no interest for each flagged volume (scrubbing).

In addition, functional data were denoised using a standard denoising pipeline (Nieto-Castanon, 2020) in CONN (Whitfield-Gabrieli & Nieto-Castanon, 2012) release 22.v2407 (Nieto-Castanon & Whitfield-Gabrieli, 2022) and SPM (Penny et al., 2007) release 12.7771 including the regression of potential confounding effects characterized by white matter time-series (5 CompCor noise components), CSF time-series (5 CompCor noise components), movement regressors and their first order derivatives (12 components), session and task effects and their first order derivatives (6 factors), and linear trends (2 factors) within each functional run, followed by bandpass frequency filtering of the BOLD time-series (Hallquist et al., 2013) between 0.008 Hz and 0.09 Hz. CompCor (Behzadi et al., 2007; Chai et al., 2012) noise components within white matter and CSF were estimated by computing the average BOLD signal as well as the largest principal components orthogonal to the BOLD average within each subject’s eroded segmentation masks. The 10 CompCor noise regressors for white matter and CSF were then added as regressors of no interest to the first-level GLMs.

### 2.4 Computational modelling of values, RPEs, and learning rate

For the learning process, we implemented the trial-wise updating mechanism using a Rescorla-Wagner (RW) model which is an established model for reinforcement learning (Rescorla & Wagner, 1972). To model basic multisensory processing involved in constructing the computational quantities of the mental model and the choice processes that translate them into responses, a drift diffusion model (DDM; Ratcliff, 1978) was implemented based on the approach outlined by Pedersen et al. (2017). This generative model allowed us to estimate parameters of the trial-wise learning process, inferred from the accuracy and reaction times of the responses from each trial (Pedersen et al., 2017). This approach permitted the calculation of trial-wise values for each multisensory association, drift rates, and RPEs.

The model used in this study is illustrated in **Figure 2**. The RW model is used to calculate the RPEs and update the values *V* associated with the responses to the current multisensory association presented in each trial. The RPE for a trial *t* is defined as the difference between the actual reward *R* and the expected reward (= value *V*):

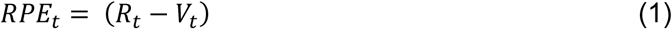

**Figure 2.**
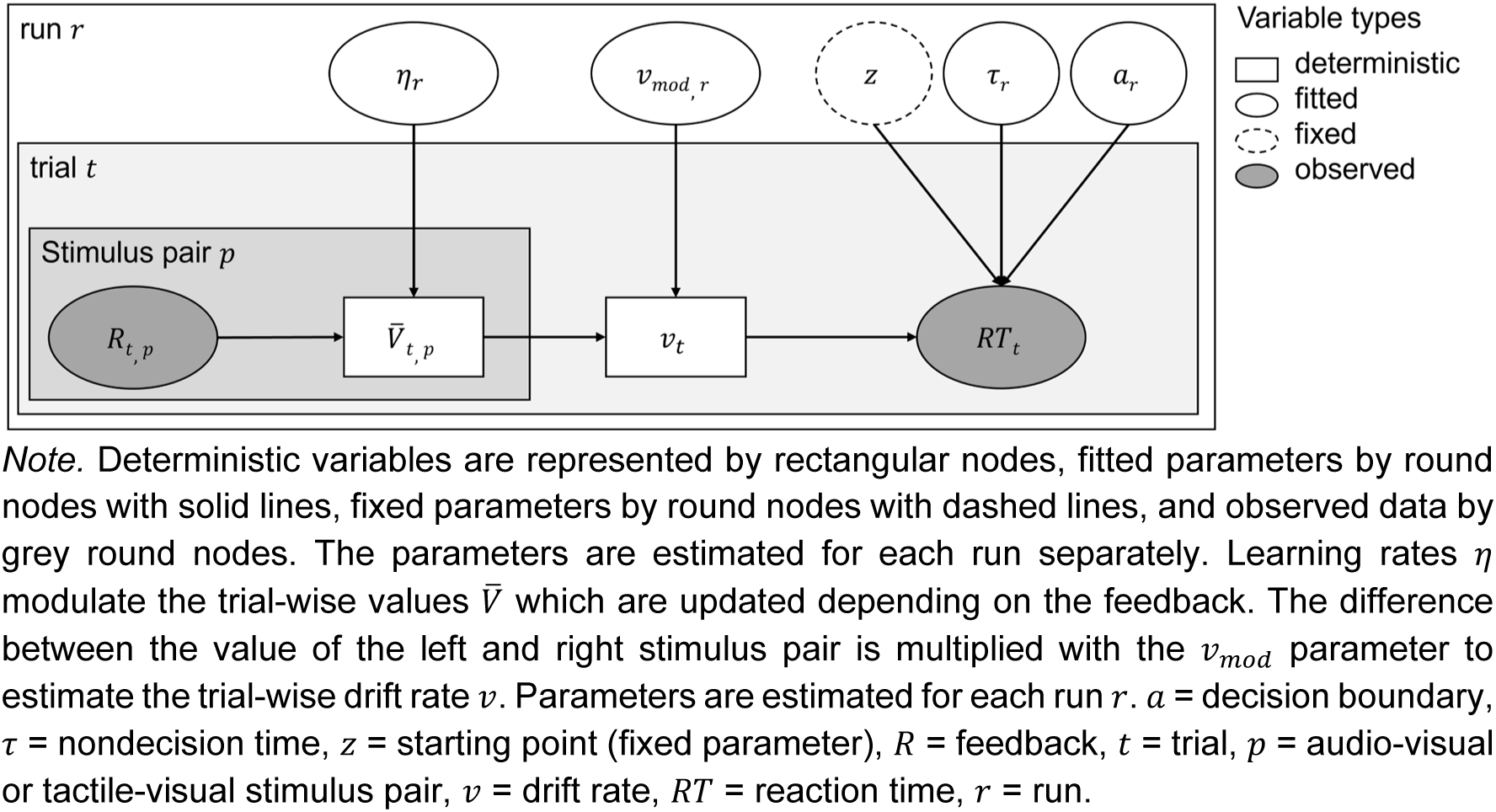
Schematic of the reinforcement learning drift diffusion model.

Then, the value *V* for each trial *t* was updated by the RPE weighted by the learning rate η:

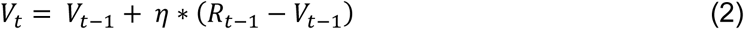

Therefore, the learning rate is a weighting factor that indicates the extent to which values are updated after each trial (for a more detailed interpretation, see Zhang et al. (2020)). In this study, only the value for the chosen response to the specific multisensory stimulus combination was updated.

Then for each trial *t*, the values for the left 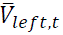 and right 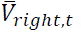 stimulus combination were normalised to ensure that the sum of both values was 1 as seen in formula 3. The value for the left stimulus *V*_left,t_ combination denotes the reward expectation for the multisensory stimulus combination of the auditory/tactile stimulus and the visual symbol displayed on the left side in trial *t*. *V*_right,t_ denotes the multisensory stimulus combination of the auditory/tactile stimulus and the visual symbol displayed on the right side in trial *t*.

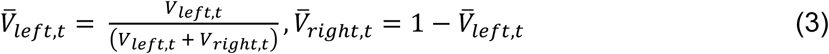

These normalised values from the RW were then entered in the DDM to calculate the drift rate *v* by multiplying the difference between the left and right value with the drift weight

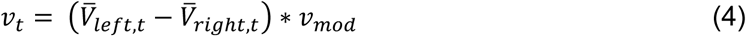

Therefore, the drift weight is a scaling factor and corresponds to the sensitivity to value differences (e.g., with a high drift weight even a small difference between the values leads to high drift rate; Pedersen et al. (2017)). Larger differences between the two expected values lead to higher drift rates and therefore faster information accumulation (e.g., when one option has a clearly higher reward expectation), while information accumulation is slower, when the options have comparable reward expectations. We used the wiener first passage time (WFPT) algorithm (Navarro & Fuss, 2009) to fit the choice behaviour and reaction time data:

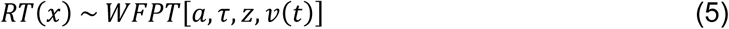

With this algorithm, the probability density for hitting the upper boundary (left choice) or lower boundary (right choice) at any given time point *t* could be calculated:

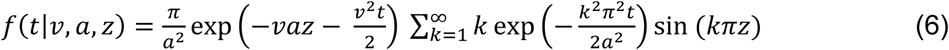

This returns the probability of choosing *x* given the observed reaction time and the expected reward. For further details, please refer to the paper by Pedersen et al. (2017).

In our approach, the learning rate η was a free parameter whereas the initial values for all stimulus combinations were fixed to 0.5 (uniform unbiased initial values). The DDM included three additional free parameters: the non-decision time τ, the boundary separation *a*, and the drift weight *v*_mod_. The starting point *Z* in the DDM was fixed to 0.5 ∗ *a*, indicating no bias towards any of the decisions. The boundary separation indicates the amount of information required until a decision criterion is reached. The non-decision time represents the amount of time that is irrelevant for the decision-making process, such as the time required for response execution. The drift rate *v* indicates the rate of accumulation of information. In the approach used in this study, free parameters (learning rate, boundary separation, non-decision time, and drift weight) were estimated separately for each run of 44 trials. The estimated parameters were then used to obtain trial-wise values for the RPE, values, and drift rate, which were used to capture the dynamic nature of learning and decision making.

Prior to applying the model to the dataset, we conducted simulations to verify that all model parameters could be recovered successfully (see Supplementary Material A.5). Then, the validated model was used for parameter estimation on the collected behavioural data and the estimated parameters were then used to identify corresponding neural processes (Farrell & Lewandowsky, 2018; R. C. Wilson & Collins, 2019). This approach ensured that our models accurately captured the underlying cognitive processes of multisensory learning. The model fitting pipeline was set up in MATLAB using the Genetic Algorithm (ga) from the Global Optimization Toolbox, in accordance with the recommendations set forth by R. C. Wilson & Collins (2019).

### 2.5 Statistical analyses

#### 2.5.1 Analyses of task performance and computational parameters

As detailed in the supplementary materials (A.6), preprocessing of the behavioural data involved the exclusion of trials with omissions, and reaction times that were either too fast or too slow, resulting in a final set of 7,594 valid trials for analysis.

To evaluate learning outcomes and differences between modalities, mean reaction times for correct trials, mean accuracy scores (correct responses divided by the total number of trials not counting omissions), and mean absolute drift rate were calculated for each bin and modality, where one bin represented one-fourth of the task (11 trials). Reaction times, accuracies, learning rate, non-decision time, boundary separation, and the absolute drift rate were then analysed using linear mixed models (library ‘lmerTest’, Kuznetsova et al., 2017) with subject as a random effect and modality and age as fixed effects. The model for the reaction times, accuracies, and absolute drift rate also included bin as fixed effect. Post-hoc analyses for significant main effects with more than one level were computed (library ‘emmeans’; Lenth, 2024) and p-values were adjusted for multiple comparisons by the Tukey method. Analyses were conducted using the R Statistical language (version 4.4.1; R Core Team, 2024). Effect sizes (η^2^) are interpreted using the following guidelines: values < .01 are considered very small, values from .01 up to (but not including) .06 are small, values from .06 up to (but not including) .14 are medium, and values > .14 are large (Field, 2013).

#### 2.5.2 Whole-brain analysis

For the first-level analysis, we built a general linear model (GLM) using the individual onsets of two regressors of interest (stimulus, feedback) and the two parametric modulators for value and RPE. The value modulator was added on the stimulus regressor and included the normalised value for the chosen stimulus combination. The RPE modulator was added on the feedback regressor and included the trial-wise RPE values. We added 4 additional regressors of no interest: 1) stimulus onset for RT outlier trials, 2) feedback onset for RT outlier trials, 3) stimulus onset for omission trials, and 4) feedback onset for omission trials to the GLM. The GLM also included six realignment parameters and one additional regressor of no interest for each flagged scan, whenever available. This was done to account for movement effects (scrubbing). The GLM included all available AV and TV runs for each subject and was convolved with the canonical hemodynamic response function implemented in SPM12.

Whole-brain second-level one-sample t-tests investigated the activation during stimulus processing, feedback processing, value processing, and RPE processing. Additionally, age was added as an effect of interest to all contrasts. For the contrasts investigating stimulus processing, the learning rate was added as an additional effect of interest (if more than one run was included, the mean learning rate from all included runs was used). Further, to investigate main and interaction effects between age and modality (AV and TV), a mixed-effects ANCOVA was conducted. The design matrix included the scans for each condition and participant. Age, condition, and their interaction were included as regressors of interest, where age is a continuous and condition a factorial variable. The model accounted for variability within subjects by including one regressor per subject. These analyses additionally included the covariates sex and handedness to control for potential confounds. The effect of age, learning rate, and the described contrasts were used to investigate general brain activity during multisensory learning and how the activation differed between modalities (AV and TV) and in relation to age. Statistical parametric maps were thresholded at a voxel-wise *p* < .001 (uncorrected), with clusters corrected for multiple comparisons using a family-wise error (FWE) rate of *p* < .05, unless stated otherwise. With this threshold, the likelihood of Type I error rates in parametric cluster correction can be reduced to an acceptable level (see Eklund et al. (2016)).

#### 2.5.3 Region of interest analyses

For region of interest (ROI) analyses, the same first-level outputs were used as for the whole-brain analyses. Masks for multisensory regions, value processing and RPE processing regions were created using neurosynth (www.neurosynth.org). We downloaded the uniformity test activation mask for each topic and then processed each mask to extract all the subclusters as individual masks (a detailed description of the processing can be found in the Supplementary Material A.7). With the code provided by MarsBaR (https://marsbar-toolbox.github.io/faq.html), we extracted the beta values corrected for the mean signal in each ROI (Pernet, 2014). We extracted the mean beta values over all subclusters and for each subcluster separately from each mask. The “multisensory network” mask was used with the AV stimulus and TV stimulus contrasts, the “value network” mask with the AV value and TV value contrasts, and the “reward prediction error network” mask with the AV RPE and TV RPE contrasts. In a next step, these beta values were then analysed using linear mixed models in R to investigate differences between modalities and the effect of age. First, we looked at the effects of modality and age and their interaction on the whole mask (beta value ∼ modality × age). Then we applied the same linear mixed model on each subcluster. In the multisensory network ROI, the learning rate was included as an additional main effect of interest. Age and learning rate were centred to the mean and the modality factor was orthogonalized. To correct for multiple comparisons, all the p-values from each linear mixed model were extracted and corrected using the False Discovery Rate (FDR) procedure (Benjamini & Hochberg, 1995). The FDR correction was applied with a significance level of α = 0.05. This allowed us to control for the proportion of false positives among the test results.

## 3 Results

### 3.1 Behavioural results

*General Task Performance:* the mean reaction times for AV and TV runs were 2.26s (*SD* = 0.37s) and 2.49s (*SD* = 0.35s), respectively. The median accuracy for AV and TV runs over all bins were 65.91% (*IQR* = 24.70%) and 65.12% (*IQR* = 23.58%), respectively. Bin-level accuracy inspection revealed that, with the exception of two participants, all individuals exceeded the chance threshold (≥ 63.64%) in at least two bins across all runs. Further accuracy at the end of the task (bin 4), showed a median of 72.73% correct for both AV and TV runs. As expected in a learning task, these results clearly exceeded chance level. To enable a valid comparison of learning across modalities, we did not exclude participants, who surpassed chance performance in only one of the two modalities. An overview including omissions and outliers can be found in the supplementary material B.2.

Linear mixed models were computed to examine the effect of sensory modality (AV vs TV), bin (1, 2, 3, 4) and age on reaction times and accuracy.

*Reaction Time:* Residuals from the linear mixed-effects model on raw RT showed no significant deviation from normality (DHARMa KS *p* = .105; dispersion *p* = .856), whereas the log-transformed model exhibited significant quantile and outlier deviations (DHARMa KS *p* < .001; dispersion *p* = .856). Accordingly, all inferential statistics reported below are based on the untransformed RT model. The best fitting LMM for reaction times (reaction time ∼ modality + bin + age + modality:age + (1 | participant)) showed no significant main effect of modality (F(1,464) = .06, *p* = .814; η^2^ = .0001, 95% CI [.00, 1.00]), but a significant main effect of bin (F(3,464) = 8.83, *p* < .001; η^2^ = .05, 95% CI [.02, 1.00]), and age (F(1,65) = 10.66, *p* = .002; η^2^ = .14, 95% CI [.04, 1.00]), and a significant interaction between modality and age (F(1,464) = 5.59, *p* = .018; η^2^ = .01, 95% CI [.001, 1.00]). Adding the interactions between modality and bin, age and bin, and the three-fold interaction between modality, age, and bin did not significantly improve model fit (*p*s > .0854). Post-hoc pairwise comparisons between the bins can be seen in Table B2.2 in the Supplementary Material. The results show significantly faster RTs in the third and last bin compared to the first and second bin. RTs were also faster the older the children, but this effect was weaker in TV compared to AV runs (see **Figure 3A** and **C**).

**Figure 3.**
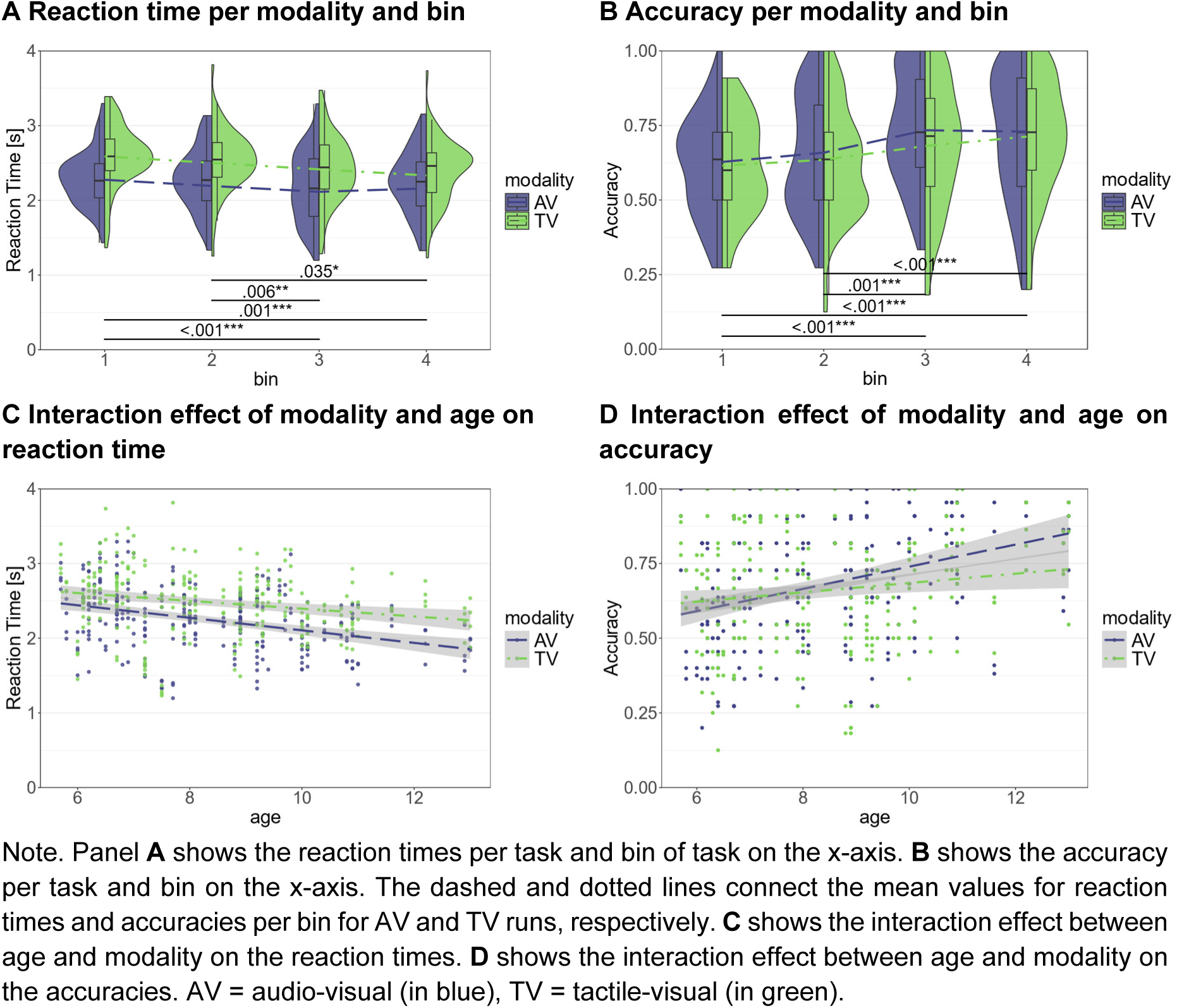
Reaction time and accuracy per modality and task bin.

*Accuracy:* For the accuracy, the best fitting LMM (accuracy ∼ modality + bin + age + modality:age + (1 | participant)) showed significant main effects of modality (F(1,464) = 6.70, *p* = .010; η^2^_p_ = .01, 95% CI [.002, 1.00]) at the intercept (age = 0), bin (F(3,464) = 13.85, *p* < .001; η^2^_p_ = .08, 95% CI [.04, 1.00]), and age (F(1,65) = 10.51, *p* = .002; η^2^p = .14, 95% CI [.03, 1.00]), and a significant interaction between modality and age (F(1,464) = 8.79, *p* = .003; η^2^_p_ = .02, 95% CI [.004, 1.00]). Adding the interactions between modality and bin, age and bin, and the three-fold interaction between modality, age, and bin did not significantly improve model fit (*p*s > .163). Post-hoc pairwise comparisons between the bins can be seen in Table B2.2 in the Supplementary Material. The results show significantly higher accuracies in the third and last bin compared to the first and second bin, and with increasing age of the children, although this effect is less strong in TV runs compared to AV runs (see **Figure 3B** and **D**). Although the uncentered model estimated a modality effect at the intercept (age = 0), this effect became non-significant when age is mean centred prior to fitting the LMM, while all other effects remained unchanged.

### 3.2 Modelling results

Linear mixed models were conducted to examine the effect of modality (AV vs. TV) and age on the learning rate, the non-decision time, and the boundary separation. For the drift rate, we fitted the same model as for the reaction time and accuracy, with modality (AV vs. TV), bin (1, 2, 3, 4), and age as predictors.

*Learning rate:* Mean learning rates were 0.19 (*SD* = 0.26) in AV and 0.20 (*SD* = 0.26) in TV runs. The best fitting LMM included only an intercept (and random intercepts for participants) which was significantly greater than zero (*β* = 0.20, 95% CI [.15, .24], *t*(131) = 8.34, *p* < .001). Adding the main effect of either modality and age, or the interaction between the two, did not improve model fit (*p*s > .271). These results indicate that learning rates were significantly different from 0, but did not vary with age or differ between AV and TV learning.

*Non-decision time*: Mean non-decision times were 0.98 (*SD* = 0.40) in AV and 1.14 (*SD* = 0.44) in TV runs. The best fitting LMM (non-decision time ∼ modality + age + (1 | ID)) showed a significant main effect of modality (F(1,66) = 9.61, *p* = .003; η^2^_p_ = .13, 95% CI [.03, 1.00]) and age on (F(1,65) = 6.27, *p* = .015; η^2^_p_ = .09, 95% CI [.01, 1.00]) the non-decision time. These results indicate significantly longer non-decision times and thus prolonged multisensory processing in TV compared to AV runs (see **Figure 4A**), and shorter non-decision times with increasing age (see **Figure 4B**). Adding the interaction between modality and age to the model, did not improve model fit (χ^2^(1) = 0.66, *p* = .416), indicating no significant interaction effect.

**Figure 4.**
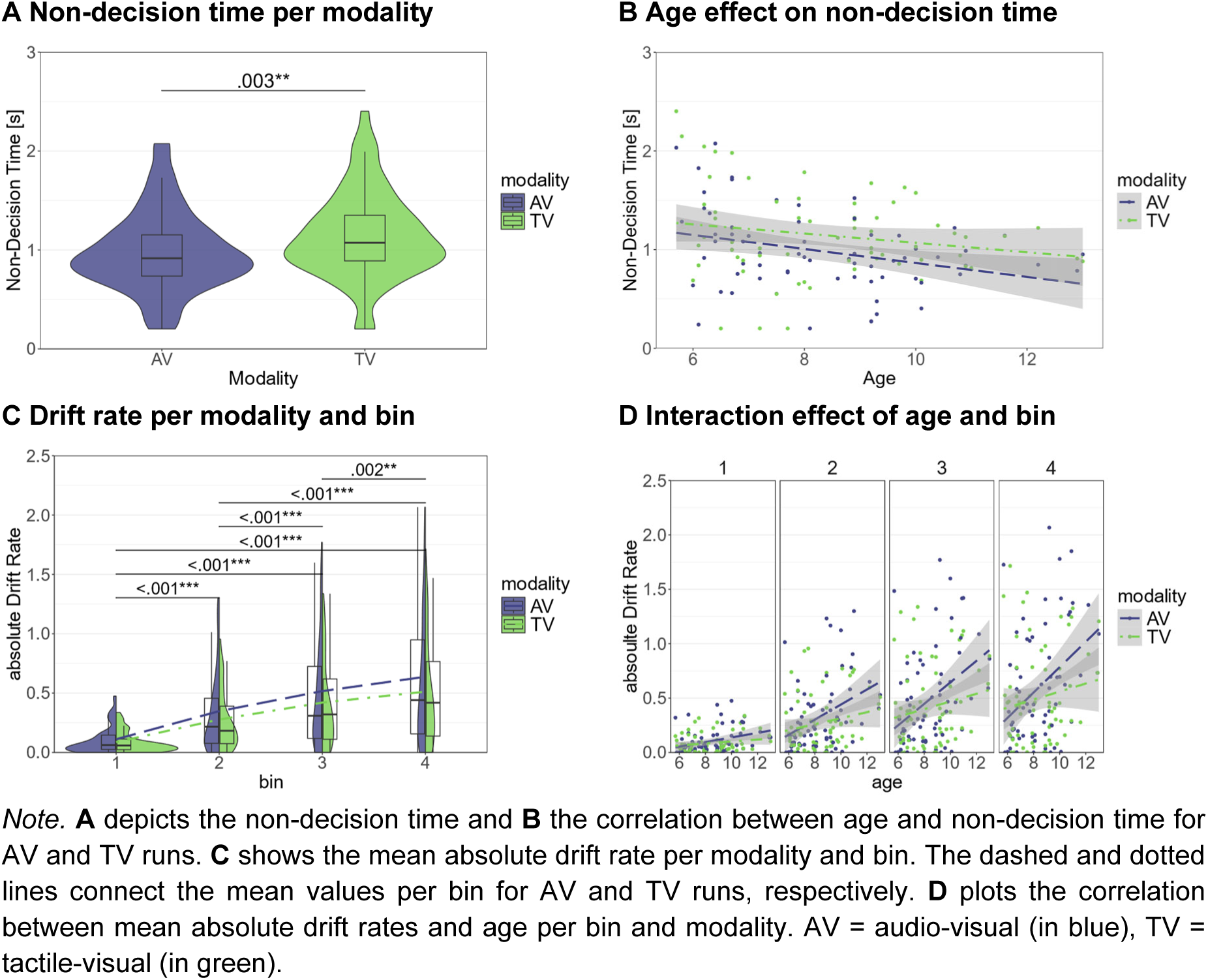
Modality and age effects on non-decision time, boundary separation, and drift rate.

*Boundary separation:* Mean boundary separation values were 2.46 (*SD* = 0.35) in AV and 2.54 (*SD* = 0.41) in TV runs. The best fitting LMM included only an intercept (and random intercepts for participants) which was significantly greater than zero (*β* = 2.5, 95% CI [2.43, 2.57], t(131) = 67.44, *p* < .001). Adding the main effect of either modality and age, or the interaction between the two, did not improve model fit (*p*s > .188). These results indicate that boundary separations were significantly different from 0, but did not vary with age or differ between AV and TV learning.

*Drift rate:* Mean absolute drift rate values for the 1^st^, 2^nd^, 3^rd^, and last bin were 0.10 (*SD* = 0.11), 0.33 (*SD* = 0.33), 0.48 (*SD* = 0.45), 0.59 (*SD* = 0.53), respectively, in AV and 0.09 (*SD* = 0.09), 0.27 (*SD* = 0.25), 0.41 (*SD* = 0.37), 0.50 (*SD* = 0.44), respectively, in TV runs. Results of the best fitting LMM on the drift rate (absolute drift rate ∼ modality + bin + age + modality:age + bin:age + (1 | participant)) show a significant main effect of modality (F(1, 461) = 14.57, *p* < .001; η^2^ = .03, 95% CI [.01, 1.00]) and age (F(1,65) = 8.29, *p* = .005; η^2^ = .11, 95% CI [.02, 1.00]), but not bin (F(3,461) = 0.17, *p* = .920; η^2^ = .001, 95% CI [0.00, 1.00]). There was also a significant interaction between modality and age (F(1,461) = 20.84, *p* < .001; η^2^ = .04, 95% CI [.02, 1.00]) and between bin and age (F(3,461) = 6.32, *p* < .001; η^2^ = .04, 95% CI [.01, 1.00]). This indicates higher absolute drift rates and thus faster evidence accumulation and more efficient decision making in older children, but this age effect on decision efficiency is stronger in AV compared to TV runs and in later compared to earlier bins (see **Figure 4C** and **D**). Post-hoc pairwise comparisons between the bins can be seen in **Table 2**.

**Table 2.** Post-hoc tests of the effect of bin on drift rate.

| Model | contrast | estimate | SE | df | t | p |
| --- | --- | --- | --- | --- | --- | --- |
| Drift rate ~ bin | 1st - 2nd | -0.20 | 0.03 | 461.00 | -7.36 | < .001*** |
|  | 1st - 3rd | -0.35 | 0.03 | 461.00 | -12.72 | < .001*** |
|  | 1st - 4th | -0.45 | 0.03 | 461.00 | -16.33 | < .001*** |
|  | 2nd - 3rd | -0.15 | 0.03 | 461.00 | -5.36 | < .001*** |
|  | 2nd - 4th | -0.25 | 0.03 | 461.00 | -8.97 | < .001*** |
|  | 3rd - 4th | -0.10 | 0.03 | 461.00 | -3.61 | .002** |

*Note.* Post-hoc pairwise comparisons for the effect of the specified factor on the dependent variable. Estimates represent the mean difference between conditions. SE = standard error, df = degrees of freedom, p values are Bonferroni-corrected for multiple comparisons.

**Table 3.**
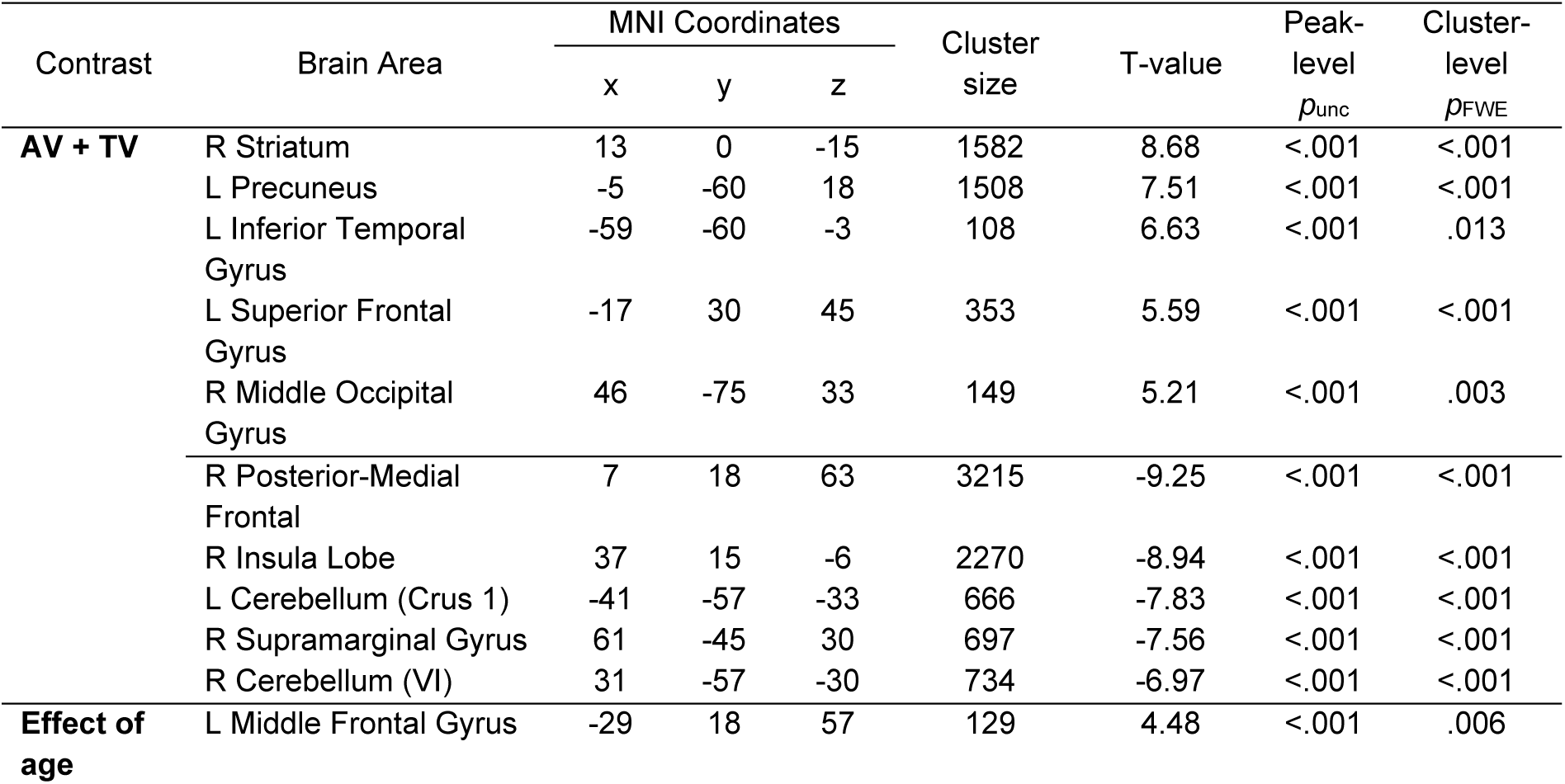
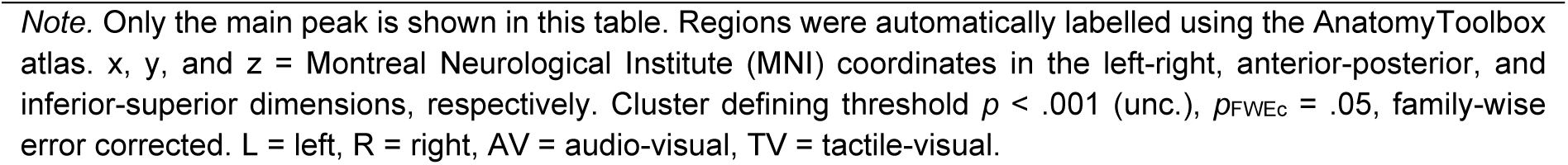
Significant clusters showing a positive and negative modulation by RPEs during feedback processing.

| Contrast | Brain Area | MNI Coordinates |  |  | Cluster size | T-value | Peak-level | Cluster-level |
| --- | --- | --- | --- | --- | --- | --- | --- | --- |
| | | x | y | z | | | $p_{unc}$ | $p_{FWE}$ |
| **AV + TV** | R Striatum | 13 | 0 | -15 | 1582 | 8.68 | <.001 | <.001 |
|  | L Precuneus | -5 | -60 | 18 | 1508 | 7.51 | <.001 | <.001 |
|  | L Inferior Temporal Gyrus | -59 | -60 | -3 | 108 | 6.63 | <.001 | .013 |
|  | L Superior Frontal Gyrus | -17 | 30 | 45 | 353 | 5.59 | <.001 | <.001 |
|  | R Middle Occipital Gyrus | 46 | -75 | 33 | 149 | 5.21 | <.001 | .003 |
|  | R Posterior-Medial Frontal | 7 | 18 | 63 | 3215 | -9.25 | <.001 | <.001 |
|  | R Insula Lobe | 37 | 15 | -6 | 2270 | -8.94 | <.001 | <.001 |
|  | L Cerebellum (Crus 1) | -41 | -57 | -33 | 666 | -7.83 | <.001 | <.001 |
|  | R Supramarginal Gyrus | 61 | -45 | 30 | 697 | -7.56 | <.001 | <.001 |
|  | R Cerebellum (VI) | 31 | -57 | -30 | 734 | -6.97 | <.001 | <.001 |
| **Effect of age** | L Middle Frontal Gyrus | -29 | 18 | 57 | 129 | 4.48 | <.001 | .006 |

*Note.* Only the main peak is shown in this table. Regions were automatically labelled using the AnatomyToolbox atlas. x, y, and z = Montreal Neurological Institute (MNI) coordinates in the left-right, anterior-posterior, and inferior-superior dimensions, respectively. Cluster defining threshold $p < .001$ (unc.), $p_{FWEc} = .05$ , family-wise error corrected. L = left, R = right, AV = audio-visual, TV = tactile-visual.

### 3.3 Whole-brain fMRI results

To investigate the effects of modality and age on brain activation, we analysed the BOLD signal during both stimulus and feedback processing using a whole-brain approach. Voxel-wise analyses allowed for an unbiased exploration of task-related activation patterns across the entire brain, without prior assumptions. Additionally, we incorporated parametric modulations during these phases to examine how variables such as value and reward prediction errors (RPEs) influence BOLD responses.

#### 3.3.1 Effects of age on stimulus and feedback processing

During AV and TV stimulus processing, age correlated positively with activation in an extended visual and parietal cluster, frontal regions, temporal regions, and the supplementary motor cortex. Brain activation in bilateral post-central, parietal regions, bilateral pre– and post-central regions, medial frontal regions, and bilateral hippocampus negatively correlated with age during AV stimulus processing. During TV stimulus processing, age correlated negatively with brain activation in temporal, hippocampal, and postcentral, parietal regions (see **Figure 5A**). The main effect of age showed visual regions and the inferior frontal gyrus was more active with increasing age of the children for both AV and TV runs. The right supramarginal gyrus was less active the older the children were (see **Figure 5B**). The interaction effect indicates that the bilateral auditory showed a stronger age effect in AV compared to TV runs. Bilateral sensory regions showed a stronger age effect in TV compared to AV runs (see **Figure 5C**). The cluster table and general brain activation during stimulus processing can be found in the Supplementary Material B.3.

**Figure 5.**
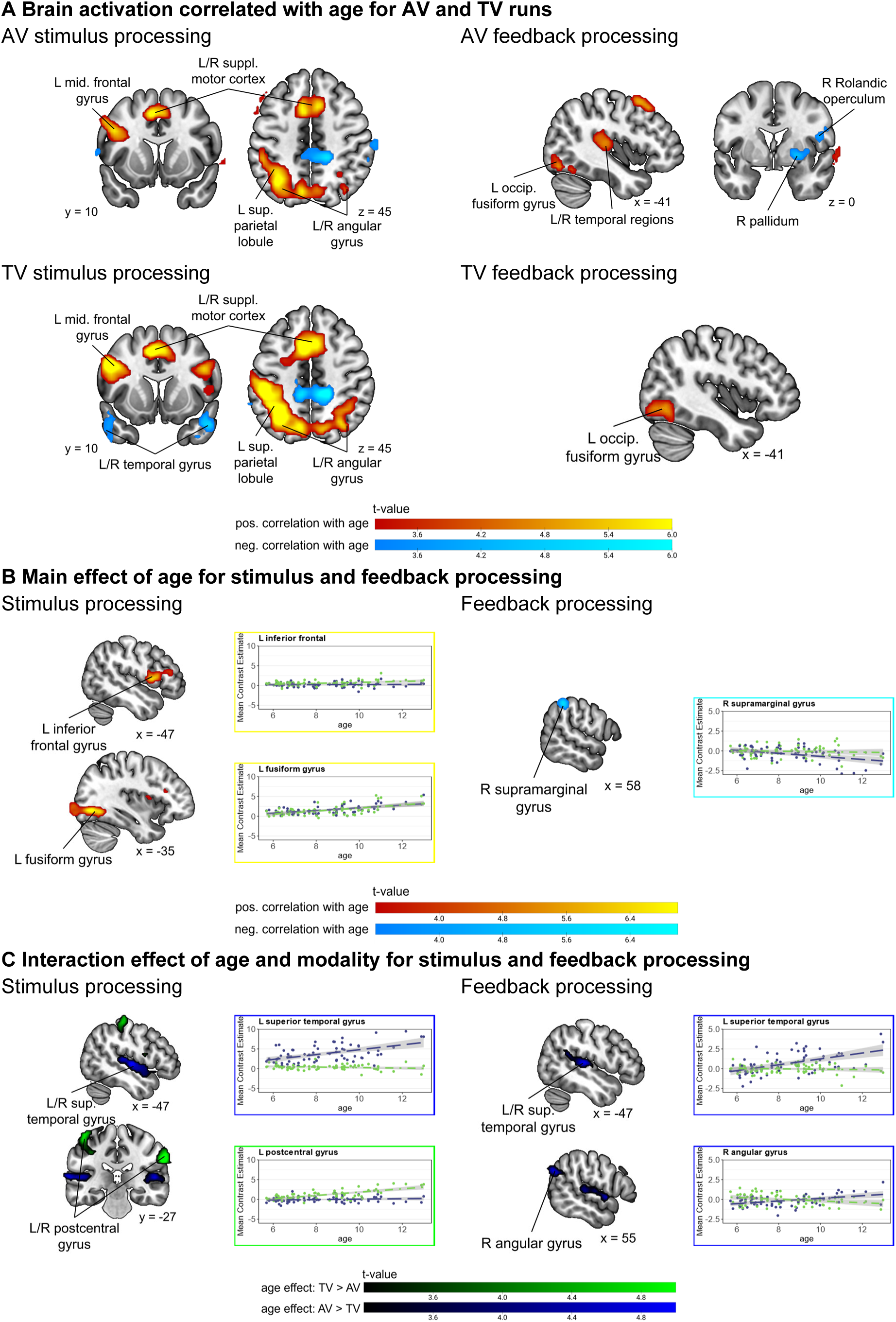
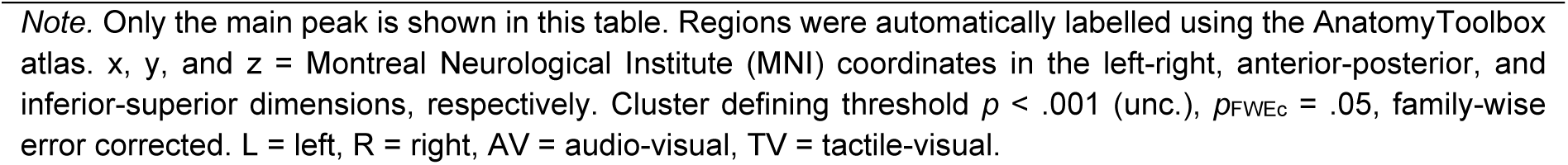
Age-effects on brain activation during stimulus and feedback processing.

**Figure 6.**
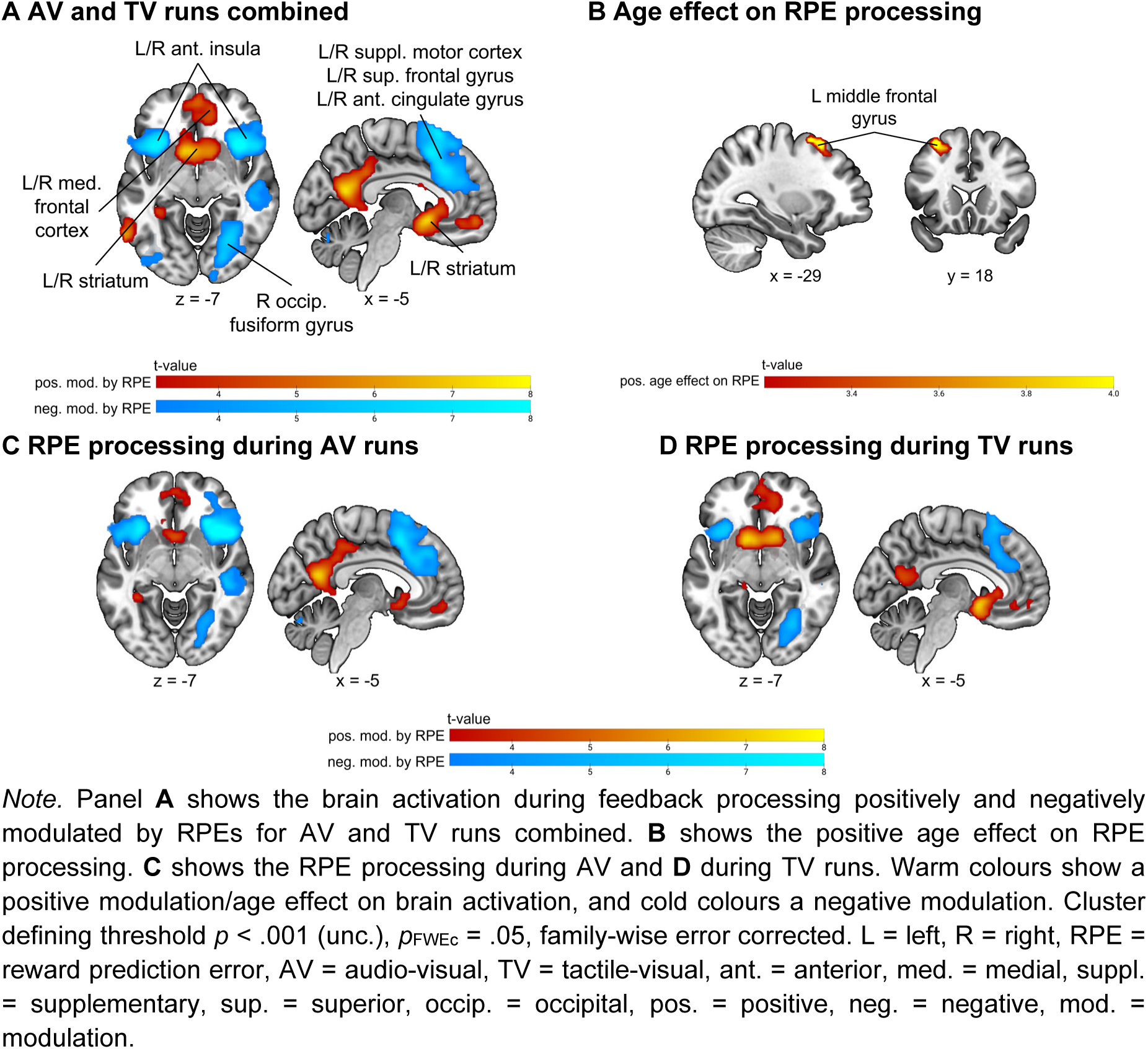
Effects of reward prediction errors on brain activation.

In AV runs during feedback processing, age positively correlated with brain activation increases in temporal, visual, and frontal regions. In TV runs during feedback processing, age correlated positively with the brain activation in visual and frontal regions (see **Figure 5A**). Brain activation in the right pallidum, left postcentral gyrus, and right Rolandic operculum showed a negative correlation with age during AV feedback processing. No significant negative correlation with age was observed during TV feedback processing. The main effect of age revealed that a cluster in the right supramarginal gyrus showed lower activation with age in both AV and TV runs. There were no significant clusters showing higher activation with age in both AV and TV runs (see **Figure 5B**). The interaction effect between age and modality revealed that auditory regions and the right angular gyrus showed higher activation with age in AV runs. There were no significant clusters showing higher activation in older children during TV runs, but not AV runs (see **Figure 5C**; the Table B4 in the Supplementary Material shows all the clusters). The general brain activation during feedback processing can be found in the Supplementary Material B.4.

#### 3.3.2 Modulation of the BOLD signal by model parameters

### 3.4 ROI analyses

To examine the effects of age and modality more closely in brain regions commonly associated with multisensory integration, reward prediction error, and value processing, we conducted ROI analyses in a-priori selected regions of the multisensory, the RPE, and the value networks. These networks and the included regions used for these analyses are shown in **Figure 7**. ROI analyses in the value network are presented in Supplementary Material B.8, as whole-brain analyses revealed no significant activation within the value mask.

**Figure 7.**
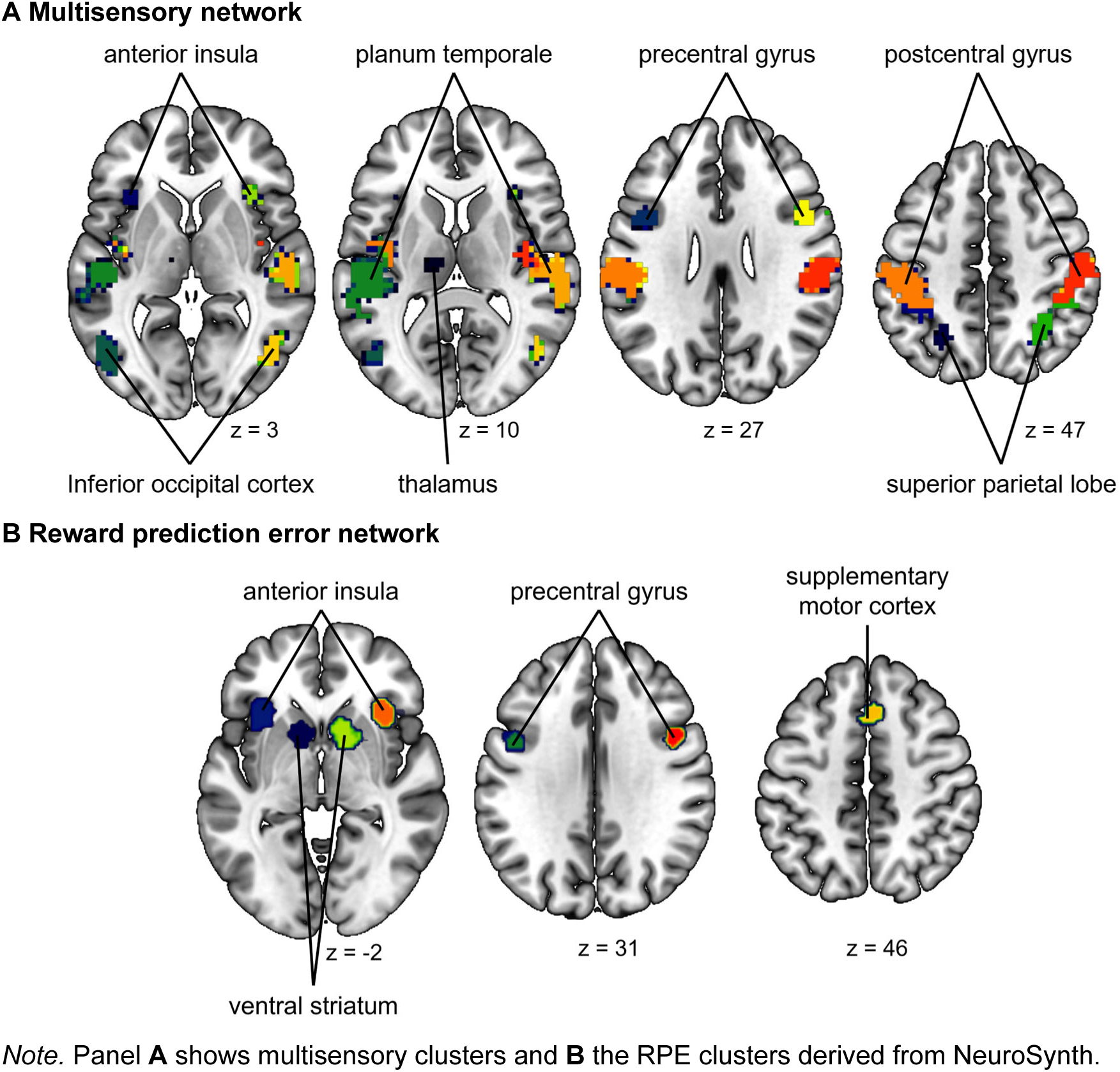
Depiction of meta-analytical masks used for ROI analyses.

#### 3.4.1 Brain activation during stimulus presentation

We first examined how brain activation during multisensory stimulus processing was affected by age and modality in regions previously associated with multisensory integration. We observed a significant main effect of modality with higher brain activation for AV runs compared to TV runs across the whole multisensory network ROI and the bilateral planum temporale. In the left postcentral gyrus, right precentral gyrus, and left superior parietal lobe brain activation was higher for TV compared to AV runs. Age positively correlated with the brain activation across the whole network ROI and in all subclusters except the right inferior occipital cortex, right superior parietal lobe and left thalamus. The effect of age was stronger in TV runs compared to AV runs in the left postcentral gyrus and the right precentral gyrus. In the bilateral planum temporale, the age effect was stronger for AV compared to TV runs (see **Figure 8**). The learning rate positively correlated with brain activation across the whole network, in the right planum temporale, and the right postcentral gyrus (the table with the effects can be found in Supplementary Material B.6).

**Figure 8.**
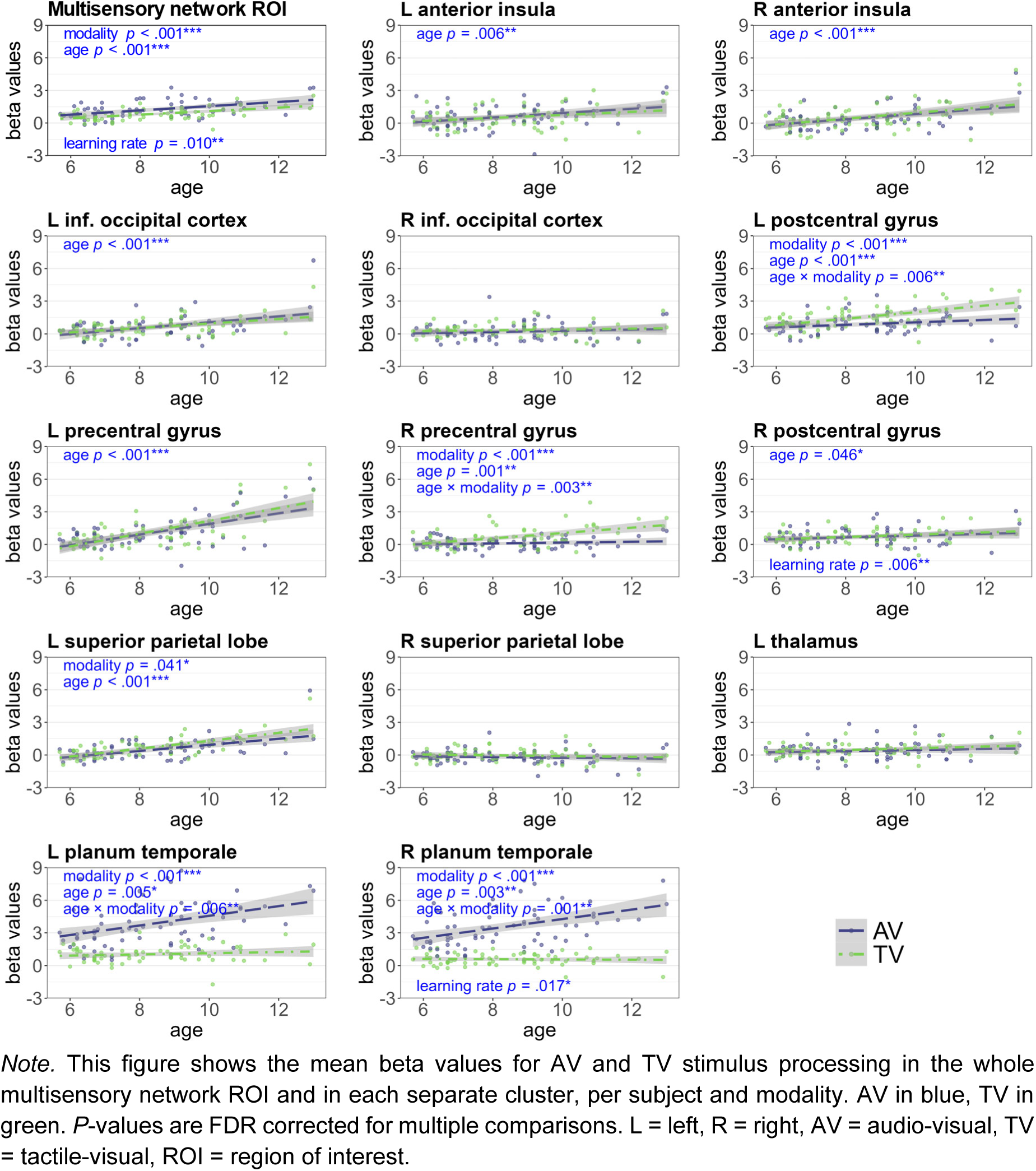
Beta values in the multisensory network ROI and distinct clusters.

#### 3.4.2 Brain activation during reward prediction error processing

Here, we examined how the parametric modulation of brain activation by trial-wise reward prediction errors was influenced by age and modality. Age negatively correlated with the brain activation across the whole RPE network, in the bilateral anterior insula, the right precentral gyrus, and the bilateral supplementary motor cortex. The negative correlation with age in several subclusters indicates higher brain activation with age to lower RPEs. The effect of age across the whole network is most likely driven by the effects in these subclusters. The brain activation across the whole network and in the right striatum shows an effect of modality, which indicates a more positive modulation by RPEs in TV compared to AV runs (see **Figure 9**; the table with model effects can be found in Supplementary Material B.7).

**Figure 9.**
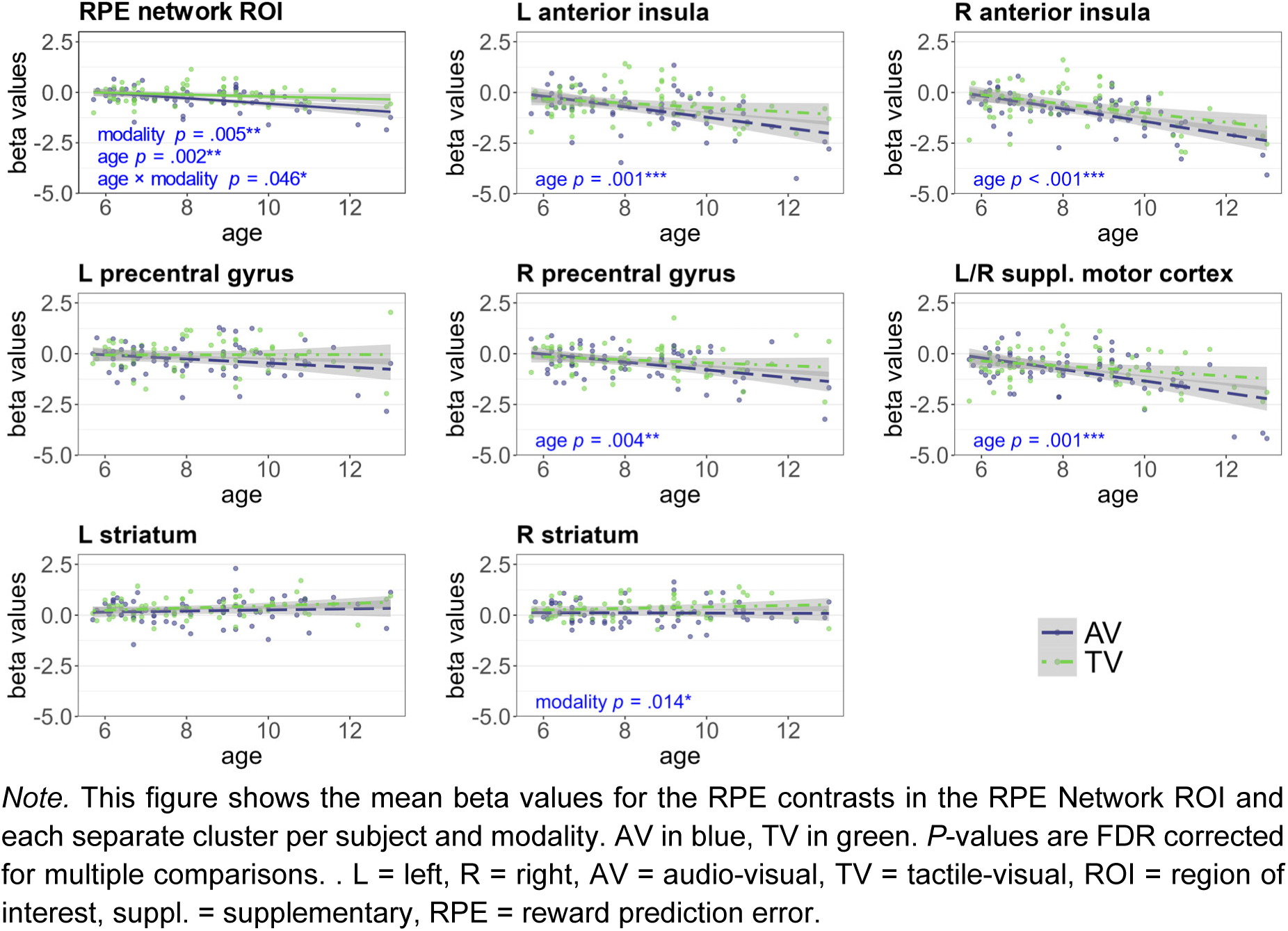
Beta values in the reward prediction error network ROI and distinct clusters.

## 4 Discussion

This study provides insights into the behavioural and neural processing underlying multisensory learning in children, focusing on audio-visual and tactile-visual associations. Three key findings emerged: First, tactile-visual associations were harder to learn than audio-visual associations as shown by performance data across ages. Second, despite considerable individual variability in learning performance, a clear developmental pattern emerged: Older-aged children demonstrated faster processing speed and greater decision-making efficiency, and this enhanced behavioural performance corresponded to higher activation in brain regions for sensory processing, cognitive control, MSI, and memory retrieval. This effect was comparable between AV and TV runs and aligns with known neural maturation processes that support multisensory learning as children develop. Third, while the core processes of reward-based learning and RPE processing, which were modality-independent, remained relatively stable across middle childhood, the older the children were, the higher their brain activity for more negative RPE, suggesting an increased sensitivity to negative feedback. This may indicate a more refined approach to evaluating rewards and punishments with development and suggests a developmental shift in how children process and utilize feedback information to guide their learning.

These findings reveal modality– and age-related alterations in behavioural and neural processes during multisensory learning, highlighting stable core learning processes and developmental shifts in neural engagement. They further emphasize the importance of accounting for age and modality in MSI research.

### 4.1 Modality-specific processing and learning across middle childhood

Behavioural analyses revealed significant learning during the multisensory task across both modalities. This improvement was shown by faster reaction times and higher accuracies in the second half of each experimental run. We only observed distinct differences in performance between the two sensory modalities in the reaction time. Tactile-visual runs consistently showed slower reaction times compared to audio-visual runs, indicating that tactile-visual multisensory processing was prolonged compared to audio-visual multisensory processing. This modality-specific difference in learning difficulty became more pronounced with increasing age. Computational modelling provided deeper insights into the learning and decision-making in multisensory learning, clarifying modality-specific and age-related changes. First, we discuss how the sensory modality influence learning processes and second how age affects learning across modalities.

The learning rate, which reflects the influence of RPEs on value updating (Zhang et al., 2020), showed similar updating of reward expectation regardless of sensory modality. Similarly, the boundary separation did not differ between modalities, indicating similar amount of information needed until a decision can be reached. However, other computational parameters differed significantly between modalities. Specifically, non-decision time was longer, and drift rates were lower in TV compared to AV learning. These findings align with slower reaction times for tactile-visual association learning, suggesting a slower accumulation process for decision-making in TV runs (Friedrich & Beste, 2019; Ratcliff & McKoon, 2008).

Consistent with the response time analyses, computational modelling revealed that processing speed improved with increasing age, driven by increased drift rates. Older children had higher drift rates in both audio-visual and tactile-visual modalities, with a more pronounced increase over time in audio-visual learning, reflecting faster and more efficient information accumulation during decision processes (Ratcliff & Rouder, 1998). Non-decision time decreased with age, indicating that the additional processing demands of tactile encoding were decreasing with age. Additionally, thresholds for accumulating information did not vary with age. In summary, while learning rates were consistent across modalities and ages, modality influenced non-decision time and drift rate, with tactile-visual pairings requiring more extensive processing. In contrast, age enhanced processing speed and decision efficiency across modalities. This suggests that maturation enhances children’s ability to associate relevant sensory information, improving speed and accuracy in multisensory learning tasks (Barutchu et al., 2020). To investigate the neural underpinnings of these age-related behavioural differences, we analysed brain activation patterns during stimulus and feedback processing at the whole-brain level and in specific ROIs.

### 4.2 Age-related recruitment of higher-order brain regions during multisensory learning

Whole-brain ANCOVA analyses showed a positive correlation between age and brain activation during stimulus processing in visual regions and the inferior frontal gyrus. This suggests that older children may engage in more detailed visual analyses to integrate visual, auditory, and tactile information during stimulus processing. The inferior frontal gyrus has been linked to working memory, episodic memory, and attention (Boisgueheneuc et al., 2006; Diveica et al., 2023; Li et al., 2013), indicating that older children engage these processes more strongly during stimulus processing independent of modality. While younger children also recruit higher-order regions for MSI, they may do so less efficiently than older children, highlighting ongoing MSI network maturation.

Brain activation during stimulus processing increased with age in the occipital cortex, precuneus, the supplementary motor area (SMA), middle frontal gyrus, and parietal cortex. This increased neural activity reflects the increased engagement of neural resources in older children to successfully perform the demanding learning task. Enhanced intraparietal sulcus activity is linked to cross-modal integration (Sours et al., 2017), mnemonic processes during both multimodal encoding and retrieval (Tibon et al., 2019), and attention (Boeken & Markett, 2023), indicating a more focused attention on relevant visual stimuli supporting encoding and learning of multisensory associations. The angular gyrus, a cross-modal hub (Hirst et al., 2021; Seghier, 2013; Tibon et al., 2019), integrates information from multiple sensory modalities, which is crucial for forming multisensory associations. The superior parietal lobule, implicated in attention processes, supports the selection, and focus on relevant sensory information during learning (Alahmadi, 2021). Greater activity in the SMA, associated with motor planning (Chung et al., 2005) and working memory (Bahlmann et al., 2009; Chung et al., 2005; Cona & Semenza, 2017), suggests that older children may plan their responses to a greater extent which corresponds to the higher accuracies in older children. Additionally, the increased middle frontal gyrus activity, tied to working memory and cognitive control (Eichenbaum, 2017), implies a more efficient knowledge access, retrieval, and utilisation during multisensory learning. Modality-specific age-related increases were observed in the superior temporal lobe during AV stimulus processing involved in auditory perception and short-term auditory memory (Crinion et al., 2006; Joseph, 1990). This points towards a greater reliance of older children on auditory memory to consolidate and retrieve learned associations. During TV stimulus processing, modality-specific age-related increases in brain activation were observed in bilateral sensory regions. Indicating that the older children the more are task-relevant sensory regions recruited during multisensory stimulus processing and thus enhancing perception, processing, and subsequently multisensory learning. In the previous chapter we described an increased efficiency in multisensory learning with increasing age. The neural results described here align with the behavioural results, suggesting that increased recruitment of sensory regions as children grow older facilitates multisensory processing and subsequently enhances multisensory learning. This is supported by previous research that showed a positive correlation between age and BOLD activation in task-relevant brain regions in a developmental sample (Schapiro et al., 2004), enhanced recruitment of task-specific neural resources to preserve task performance when task demands increase (Wessa et al., 2013), and increased activation during a working memory task in high performing adults (Nagel et al., 2009).

Modality-specific age-related decreases in activation were observed in both AV and TV conditions in a widespread network comprising the thalamus and a cluster spanning the postcentral gyrus, superior parietal lobe, and precuneus. Interestingly, younger children exhibited higher activation in the bilateral middle temporal gyrus during TV processing, suggesting increased engagement in semantic processing (Rolls, 2021). This indicates that younger children may associate semantic meanings to tactile stimuli to enhance memory and retrieval. We hypothesised that younger children would rely more on subcortical and primary sensory regions, while older children would engage higher-order cortical areas during MSL, indicating the increased employment of neural resources (Murray et al., 2016). This can partly be confirmed by the positive correlation with age during AV and TV stimulus processing in higher order brain regions such as parietal and frontal regions. We also observed negative correlations with age in subcortical regions, such as the thalamus, and primary sensory regions, such as temporal and central regions, during multisensory processing. But our analyses highlight, that the age-related changes in our sample appear to be more complex, with positive and negative correlations in both primary sensory and high-order brain regions. These findings suggest that age-related changes in neural activation patterns vary depending on task demands and the specific brain regions involved.

The results of the whole-brain analyses were corroborated by ROI analyses, focusing on regions identified as relevant for multisensory processing and integration in a Neurosynth meta-analysis. Older children exhibited greater activation in most of the regions related to MSI during stimulus processing, although in auditory regions, this increase was limited to AV processing, and in sensory regions specific to TV processing. These results show that older children engage sensory processing, cognitive control, MSI, attention allocation and memory regions more extensively. This enhanced neural engagement likely reflects greater cognitive effort, improving performance in multisensory learning tasks.

### 4.3 Differential brain activation in AV and TV feedback processing with age

On a whole-brain level, brain activation during AV feedback processing positively correlated with age in the middle frontal gyrus. This region, anatomically connected to the superior and inferior frontal gyri, supports working memory, episodic memory, and attention (Boisgueheneuc et al., 2006; Diveica et al., 2023; Li et al., 2013). Modality specific analyses revealed a positive correlation between brain activation in auditory regions and age during visual feedback processing in AV runs, suggesting older children utilize auditory memory traces to evaluate feedback and adjust behaviour to a greater extent. Similarly, increased visual activation with age in AV and TV runs indicates an increased reliance on visual memory traces for the integration of multisensory information. The fusiform gyrus, crucial for visual attention (Chen et al., 2019; Ruiz-Rizzo et al., 2018), and letter and word form processing (Dehaene & Cohen, 2011; Thesen et al., 2012), facilitates the formation of audio-visual percepts, such as letter-speech sound associations (Karipidis et al., 2018, 2021; Pleisch et al., 2019). In contrast, age-related positive correlations during TV feedback processing were limited to visual regions, likely due to the increased difficulty of integrating tactile information, which limited age-related effects. This aligns with lower TV performance and suggests similar feedback processing and learning strategies across ages. The fMRI results further align with the drift rate findings, showing more pronounced age-related changes in the AV than TV learning.

### 4.4 Modality-independent reward prediction error processing

Our findings revealed that that RPE processing was largely modality-independent on the whole-brain level, with RPEs positively modulating brain activity in the ventral striatum, known for encoding RPEs, as well as the medial frontal cortex and hippocampus, linked to learning and memory (Cohen & Ranganath, 2005; Fraga-González et al., 2025; Garrison et al., 2013; McClure et al., 2003; O’Doherty et al., 2003; Pagnoni et al., 2002). RPEs negatively modulated brain activation in regions typically associated with negative RPE signalling, such as the cingulate cortex, anterior insula, and frontal areas (Fouragnan et al., 2018). ROI analyses further confirmed the modality independent RPE processing. Age-related effects on RPE processing were found in the middle frontal gyrus in whole-brain analyses, which indicates an increased involvement of working memory, episodic memory, and attention (Boisgueheneuc et al., 2006; Diveica et al., 2023; Li et al., 2013) with development during RPE processing. ROI analyses revealed a negative correlation between age and brain activity in the bilateral anterior insula, right precentral gyrus and bilateral supplementary motor cortex. The findings of higher anterior insula activation with negative RPEs in older children are indeed consistent in previous findings in adolescents (Hauser et al., 2015; Yaple et al., 2020). This observation supports neurodevelopmental theories proposing that the anterior insula, as an immaturely connected cognitive-emotional hub in adolescence (Smith et al., 2014), biases decision-making towards affectively influenced choices, consistent with the concept of a dominant social-affective system in adolescent cognition (Crone & Dahl, 2012; Icenogle & Cauffman, 2021; Loureiro, 2020; Smith et al., 2014; Van Leijenhorst et al., 2010). For our MSL paradigm the findings thus suggests that older children exhibit heightened sensitivity to negative or unexpected feedback, processing it more extensively, thereby facilitating more effective learning adjustments and error correction. But our findings indicate that this sensitivity is not limited to the anterior insula but extends to other RPE regions as well. In summary, as development progresses children exhibit enhanced MSI, relying on a broader network of brain regions, particularly during audio-visual learning. While RPE processing is modality-independent and stable across ages, older children demonstrate increased sensitivity to negative feedback, suggesting more efficient learning from reward and punishment.

### 4.5 Limitations

While our study sheds light on how multisensory processing and learning evolve during middle childhood several limitations should be considered when interpreting these results.

First, individual differences in learning abilities, attention and motivation may have influenced performance, particularly among younger children. Eye-tracking in future studies could help assess attention and engagement more directly.

Second, the adaptive nature of the task, ensuring comparable demands across participants, may have masked subtle age-related differences in cognitive and sensory processing. The task’s inherent difficulty, especially for younger children, may have limited performance and contributed to variability in learning.

Third, although we designed the tactile vibration patterns to be distinct, they were generally more difficult to differentiate than the auditory stimuli, likely explaining the lower accuracy values during tactile-visual learning.

Further, as a cross-sectional study, our research captures a snapshot of performance at a single developmental stage. Longitudinal studies would be beneficial to track the evolution of these cognitive processes in individual children and facilitate the understanding the developmental trajectories involved in multisensory learning.

Next, we did not observe any significant correlations between brain activation and either estimated values or learning rate. This may, in part, reflect the nature of the task, which was not explicitly designed to manipulate value and may therefore lack the sensitivity needed to detect robust neural correlates of value processing. Additionally, learning rates were estimated as a single value per run and showed limited variability across participants, age, or modality (see Chapter 3.2). As a result, the between-subject variance of this parameter may have been insufficient to detect meaningful associations with brain activation.

Furthermore, although our findings indicate learning-related changes in both behaviour and model-derived parameters, and all children completed an extensive familiarisation and practice session with the tasks, we cannot entirely rule out the possibility that some observed effects may be attributable to task familiarisation rather than solely associative learning.

Further, ten participants in our sample either had a clinical diagnosis (ADHD, developmental language delay, or developmental dyslexia) and/or a clinically significant T-value on the CBCL/6-18R. Analyses without this subgroup were mostly identical for the drift rate findings, age effects on stimulus processing, neural RPE processing on the whole-brain and ROI-level. The main effect of age and the interaction effects between age and modality during feedback processing on the other hand showed some differences (see Supplementary Material B.1). While all effects, except the stronger age effect during AV feedback processing in the angular gyrus, of the whole sample could be replicated in the smaller group, there were some additional clusters in the smaller sample. Thus, these findings have to be verified in further studies.

Finally, the structure of our task is no suited to directly assess multisensory integration. Since each trial presents both a matching and a non-matching visual stimulus, it is difficult to isolate multisensory integration effects from learning effects, i.e. by contrasting congruent with incongruent multisensory pairs. A follow-up task after the multisensory learning task, specifically designed to test integration (e.g., by comparing learned matching versus non-matching combinations), would be necessary to make claims about multisensory integration of newly learned multisensory associations.

The present findings thus provide valuable insights into how multisensory processing and learning are influenced by modality and age during middle childhood. Understanding how multisensory processing and learning unfold across development lays the groundwork for studying and further understanding impairments in these processes (e.g., related to developmental language delays or other neurodevelopmental disorders).

### 4.6 Conclusion

Our findings show that typically developing children can learn new associations between abstract visual and auditory or tactile stimuli within an experimental session. Learning rates remained consistent across modalities and age groups, but modality-specific factors influenced decision-making. With increasing development, children exhibited more efficient multisensory processing and integration. This was accompanied by increased neural activation in regions linked to cognitive control, working memory, and MSI, indicating the usage of increased neural resources for MSL the older the children. Older children exhibited increased activation in the bilateral anterior insula, right precentral gyrus, and bilateral supplementary motor cortex in response to negative RPEs, suggesting a heightened sensitivity to negative feedback and a more nuanced learning process that integrates both positive and negative feedback for behavioural adaptation. In contrast, younger children in our study apparently relied more on basic learning mechanisms and seemed less able to flexibly adapt their behaviour based on negative feedback. Overall, older children use advanced cognitive strategies and neural resources in multisensory learning tasks, while younger children engage higher-order MSI regions less efficiently, reflecting ongoing dynamics and developmental maturation in multisensory learning. To conclude, our findings indicate that with increasing age children are more efficient at integrating information across different sensory modalities to aid learning of multisensory associations, which they may utilize more effectively to guide their behaviour. This ability is essential for advanced learning, problem-solving, and adaptation in dynamic environments.

## CRediT authorship contribution statement

**Nina Raduner:** conceptualisation, data curation, formal analysis, investigation, methodology, project administration, visualization, writing – original draft; **Carmen Providoli:** conceptualisation, data curation, investigation, project administration, writing – review and editing; **Sarah V. Di Pietro:** conceptualisation, data curation, investigation, project administration, supervision, writing – review and editing; **Maya Schneebeli:** methodology, supervision, writing – review and editing; **Iliana I. Karipidis:** conceptualisation, supervision, writing – review and editing; **Ella Casimiro:** conceptualisation, methodology; **Saurabh Bedi:** conceptualisation, methodology, writing – review and editing; **Michael von Rhein:** conceptualisation, funding acquisition, supervision, writing – review and editing; **Nora M. Raschle:** conceptualisation, funding acquisition, methodology, supervision, writing – review and editing; **Christian C. Ruff:** conceptualisation, funding acquisition, methodology, resources, supervision, writing – review and editing; **Silvia Brem:** conceptualisation, funding acquisition, methodology, project administration, resources, supervision, writing – review and editing.

## Declaration of competing interest

The authors report no competing financial interests or personal affiliations that may have affected the research presented in this paper.

## Declaration of generative AI and AI-assisted technologies in the writing process

During the preparation of this work the author(s) used ChatGPT, Perplexity, Gemini, and Quillbot in order improve language and readability. After using this tool/service, the author(s) reviewed and edited the content as needed and take(s) full responsibility for the content of the publication.

## Acknowledgments

We are very grateful for all the families and children for participating in this study. Further, we thank K. Bremer, A. Bussmann, N. Ehrhardt, E. Hohl, K. Gelic, S. Gubler, L. Hämmerli, S. Ismail, R. Jocham, I.P. Johnson, A. Lütolf, X. Michael, M. Peter, S. Schädler, J. Schiesser, M.A. Steuer, P. Sting, L. Strickler, C. Volk, and R. Wombacher for their help in planning, recruitment, and data collection. We thank Y. Cao for his contribution to implementing the drift diffusion model, P. Stämpfli for his advice on the MRI protocol, G.S.P. Pamplona and A. Haugg for their help in the statistical analyses and R. Borbás, P. Dimanova and S. Scatolin for their valuable input.

## Funding

This work was supported by the University Research Priority Program Adaptive Brain Circuits in Development and Learning (AdaBD) of the University of Zurich (project ChildBrainCircuits).

## Data and code availability

The data (no raw or pre-processed neuroimaging data) and code for this publication are available on https://osf.io/cmzne/ (DOI 10.17605/OSF.IO/CMZNE).

## Notes

### Competing Interest Statement

The authors have declared no competing interest.

### Summary of Updates

The paper has been reivsed after major revision during peer-review, including re-running the analyses related to the computational modelling. Specifically, we re-estimated the RLDDM parameters using an improved model with an extended parameter range for the non-decision time. Further, we additionally incorporated CompCor into the fMRI preprocessing pipeline to improve noise correction in the children's data. These changes led to minor alterations in some of the results, particularly concerning size and significance of activated clusters in the fMRI analyses, but the main findings and interpretations remain consistent. The revised analyses allowed us to more accurately estimate parameter values and to improve the interpretability and robustness of our findings. All revised sections have been updated accordingly.

