## Supplementary Material for "Growing Minds, Integrating Senses: Neural and Computational Insights into Age-related Changes in Audio-Visual and Tactile-Visual Learning in Children"

#### A Supplementary Material: Methods

##### A.1 Demographics

*Neurocognitive Assessments:* For this study, the following neuropsychological tests were included: Each participant's non-verbal IQ was estimated using the fluid reasoning index comprising of the matrix reasoning and the figure weights subtests from the Wechsler Intelligence Scale for Children – fifth edition (WISC-V; Wechsler, 2017) or the analogue subtests (matrix reasoning and picture concepts) from the Wechsler Preschool and Primary Scale of Intelligence – fourth edition (WPPSI-IV; Wechsler, 2012) for children under the age of six years. The verbal IQ was estimated by the Peabody Picture Vocabulary Test – fourth edition (PPVT-IV; Lenhard et al., 2015). Due to the high odd-even-split-half reliability (Cronbach's α = .97; Lenhard et al., 2015, p. 76), only even or odd trials were administered. We then calculated the mean IQ between the verbal and non-verbal IQ estimates for each participant. To measure sustained attention, the respective subtest from the Testbatterie zur Aufmerksamkeitsprüfung – Kinderversion „Das Schloss der Geister" (KiTAP) was used (Zimmermann et al., 2002). Each child also completed the rapid automatized naming task (RAN) and the parrots-subtest, measuring processing speed, from the Intelligence and Development Scales 2 (IDS-2) test battery (Grob & Hagmann-von Arx, 2018) to assess cognitive processing speed. Data on language and psychological development through an in-house questionnaire along with the Child Behavior Checklist (CBCL/6-18R; Döpfner et al., 2014) was provided by the parents or guardians of all but four children, for whom some information was missing. Two children were diagnosed with developmental language impairments (developmental language delay (DLD) or developmental dyslexia (DD)), and two children were diagnosed with attention deficit hyperactivity disorder (ADHD). The children who were diagnosed with ADHD were either unmedicated or continued their medical treatment as usual and were tested on days when they were taking their usual medication. A total of eight children, including the two who had been diagnosed with ADHD, exhibited a clinically significant T-value (greater than 63) for the total score on the CBCL/6-18R. In sum, ten children either had a formal clinical diagnosis (ADHD, DLD, or DD) and/or a clinically significant T-value on the CBCL/6-18R.

*Age differences between sexes:* The Welch two sample t-test testing the difference between the age of female and male children (mean of girls = 8.49 (2.0), mean of boys = 8.19 (1.7)) suggests that the effect is statistically not significant (difference = 0.29, $t(64.99) = 0.66$, $p = .509$).

**Table A1**

*Demographic characteristics of study participants split in age ranges*

| | 5.7-8.0 | 8.1-10.5 | 10.6-13.0 |
|---|---|---|---|
| | *M* (*SD*) | *M* (*SD*) | *M* (*SD*) |
| **N** | 34 | 24 | 9 |
| **Age [y]** | 6.86 (0.68) | 9.28 (0.61) | 11.56 (0.92) |
| **Sex [f:m]** | 17:17 | 14:10 | 5:4 |
| **Handedness[a] [l:a:r]** | 4:2:28 | 2:1:20 | 0:0:9 |
| **IQ (verbal and non-verbal)** | 107.9 (10.4) | 104.4 (11.6) | 107.1 (8.8) |
| **Rapid automatized naming [T-value]** | 51.62 (7.66) | 48.42 (6.34) | 48.89 (8.96) |
| **Processing speed [T-value]** | 51.12 (7.61) | 48.62 (8.21) | 50.33 (8.96) |
| **Sustained attention median RT[b] [PR]** | 52.78 (29.19) | 46.12 (32.35) | 61.67 (20.23) |

*Note*. For descriptive purposes, participants were stratified into three nonoverlapping age bins (5.7–8.0 y, 8.1–10.5 y, and 10.6–13 y). All inferential analyses, however, used age as a continuous variable and did not rely on these groupings. [a] One child's data are missing. [b] Percentiles are missing for two children due to the absence of normative values for their age. y = years, f = female, m = male, l = left, a = ambidextrous, r = right, RT = reaction time, PR = percentiles.

#### A.2 Stimuli and backgrounds

Here the different stimuli used for this study can be found. **Figure A2** depicts all the visual stimuli grouped by set and the corresponding backgrounds that were used. Additionally, we created three sets of tactile stimulations with the frequency patterns as depicted in **Figure A2**. All auditory stimuli are uploaded here: https://doi.org/10.17605/OSF.IO/CMZNE.

**Figure A2**

*Depiction of the stimuli used in this study*

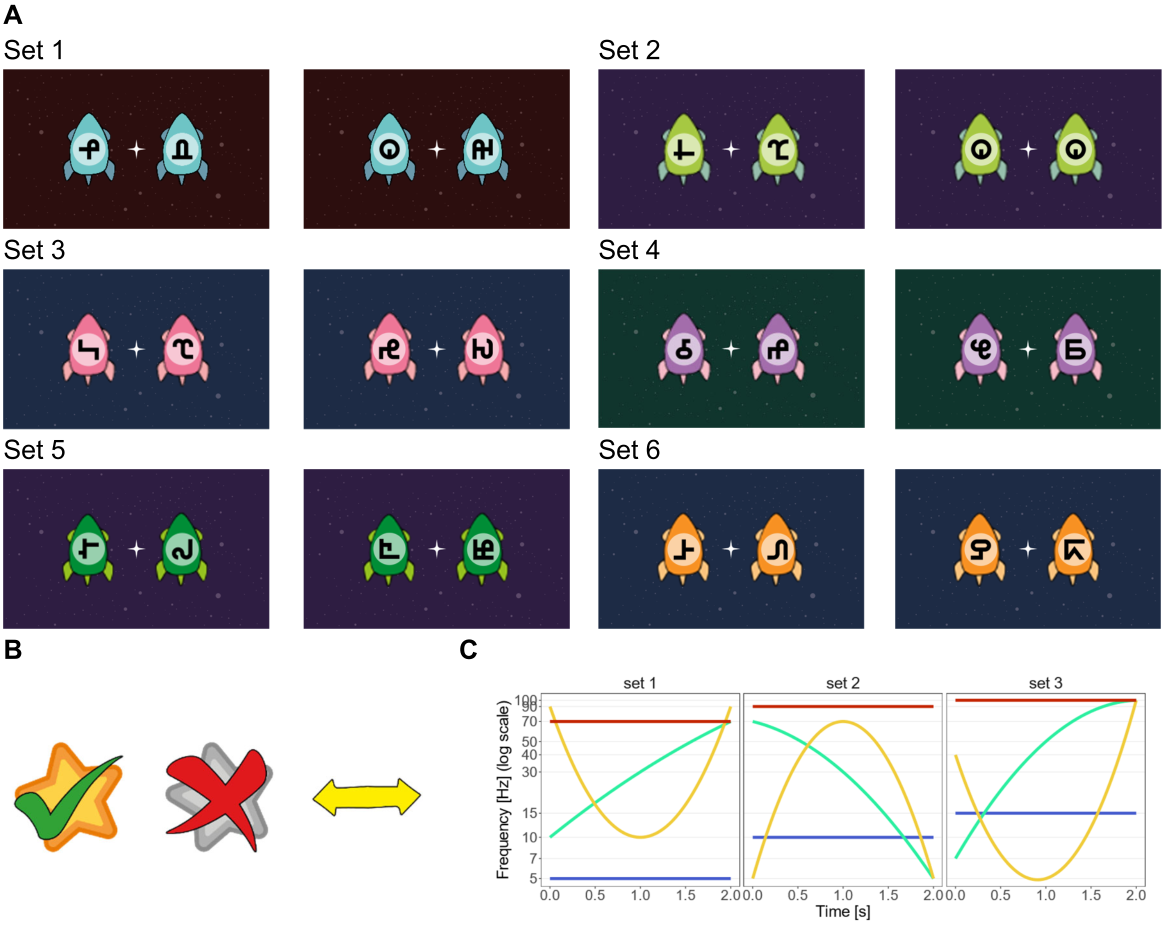


*Note.* Panel **A** shows the visual pseudo-letters used in this study and how they were grouped in sets including the corresponding backgrounds. The symbols were taken from the PseudoSloan font (Vildavski et al., 2021). **B** shows the feedback used during the multisensory learning task. From left to right: positive feedback, negative feedback, neutral feedback (when no choice was given). **C** depicts the frequency patterns for the three tactile stimulation sets. Set 1 contains a stable frequency at 5 Hz, an increasing frequency from 10 to 70 Hz, a u-shape frequency from 90 to 10 to 90 Hz, and a stable frequency at 70 Hz. Set 2 consists of a stable 90 Hz frequency, a decreasing frequency from 70 to 5 Hz, an inverted u-shape from 5 to 70 to 5 Hz, and a stable frequency at 10 Hz. Set 3 has a stable frequency at 15 Hz, an increasing frequency from 7 to 100 Hz, a u-shaped frequency from 40 to 5 to 100 Hz, and a stable frequency at 100 Hz.

#### A.3 Trial structures, probabilistic feedback, and presentation frequency

To be able to adapt the task difficulty to the individual performance, we created different trial structures. The easy trial structure allowed each auditory or tactile stimulus to be repeated once in two consecutive trials and one repetition occurred per bin of the task (e.g. auditory stimulus A appeared in trial 7 and 8). Even with the consecutive repetitions, all auditory and tactile stimuli appeared equally often within a run. In the hard trial structure, no consecutive repetition of auditory or tactile stimuli occurred. When creating the trial structures, the presentation of each auditory/tactile stimulus was balanced within 11 trials of the task to ensure that the exposure to the four matching multisensory stimulus combinations was balanced across one run. Additionally, we introduced probabilistic feedback in the task where the feedback given to the participant was wrong, e.g., negative feedback after a correct response or vice versa. There were three possible levels of difficulty: 5% probabilistic feedback (2 trials per run, one in the second and last bin), 10% probabilistic feedback (4 trials per run, one per bin of the task), or 20% probabilistic feedback (8 trials per run, two per bin of the task). The easy trial structure was always paired with 5% probabilistic feedback, the harder trial structure could be paired with all three levels of probabilistic feedback. In total, we had 8 easy and 8 hard trial structures, which were balanced within and between subjects. All pre runs were completed with the hard trial structure and no probabilistic feedback. The behavioural runs were completed with the hard trial structure and 5% probabilistic feedback. This remained unchanged for the MR session if task performance was between 60% and 80% correct. If performance was < 60%, the easy trial structure with 5% probabilistic feedback was used in the MR session. And if the performance was > 80%, the hard trial structure with 10% probabilistic feedback was used in the MR session. Thus, if accuracies were <80% in the pre-run, 5% probabilistic feedback was used in the MR run. **Table A3.1** show the distribution of all the runs across all possible difficulty levels.

**Table A3.1**

*Distribution of runs across task difficulty*

| Prob. FB | Trial structure | runs | subjs | neg. FB | pos. FB | age |
|---|---|---|---|---|---|---|
| | | *N* | *N* | *M* | *M* | *M* (*SD*) [min–max] |
| 5% | easy | 61 | 34 | 16.77 | 25.69 | 7.76 (1.57) [5.7–10.9] |
| 5% | hard | 76 | 45 | 15.33 | 27.09 | 8.99 (1.81) [5.8–13] |
| 10% | hard | 35 | 25 | 11.91 | 31.23 | 9.47 (2.11) [5.7–13] |
| 20% | hard | 8 | 7 | 17.12 | 25.75 | 10.09 (1.35) [8.1–12.2] |

*Notes.* Trial structure is coded as "hard" when no consecutive repetition of the auditory/tactile stimulus occurred within the session and as "easy" when at least one such repetition was present. Runs (*N*) denotes the number of session observations for each combination of feedback probability ("Prob. FB"; number of trials with incorrect feedback given) and task difficulty. Subjs (N) denotes the number of individual subjects for each combination. Neg./pos. FB denotes the average number of trials per session where negative or positive feedback, respectively, was given. Age is given as the mean, standard deviation, and range in years. FB = feedback, neg. = negative, pos. = positive.

In addition, the presentation frequency of the occurring stimulus combinations was manipulated within runs such that some triplets (combinations of two visual and one auditory/tactile stimulus) appeared more often than others. Each visual, auditory, or tactile stimulus appeared equally often (22/44 trials for visual stimuli, 11/44 trials for auditory/tactile stimuli) but the combinations between the modalities was manipulated. The distributions are displayed in **Table A3.2**. The combinations paired 100% indicate the matching combinations (A0, B1, C2, D3). This means that if auditory/tactile stimulus A was presented in a trial, the visual stimulus 0 was always present. But in 63.3% (7/11) of the trials, where stimulus A was presented, visual stimulus 1 was the second visual stimulus, visual stimulus 2 in 27.3% (3/11) trials, and visual stimulus 3 in 9.1% (1/11) of trials where stimulus A was presented.

**Table A3.2**

*Presentation frequency of stimulus combinations*

| *Auditory/Tactile* | **Visual** | **0** | **1** | **2** | **3** |
|---|---|---|---|---|---|
| *A* | | 100 | 63.6 | 27.3 | 9.1 |
| *B* | | 63.6 | 100 | 9.1 | 27.3 |
| *C* | | 27.3 | 9.1 | 100 | 63.6 |
| *D* | | 9.1 | 27.3 | 63.6 | 100 |

*Notes.* In this table the presentation frequency of the stimulus combinations can be seen. Auditory/tactile stimulus A was always presented with visual stimulus 0. In 63.6% of the trials, where auditory/tactile stimulus A was presented, visual stimulus 1 was the second visual stimulus. In 27.3% it was visual stimulus 2.

#### A.4 Motion during fMRI scans and age

We did not observe any effect of age on movement during MRI scans (r = 0.13, t(65) = 1.06, p = .293). For this, we correlated the mean framewise displacement (FWD) value per subject with the age of the subject (see **Figure A4**).

**Figure A4**

*Mean framewise displacement depending on age*

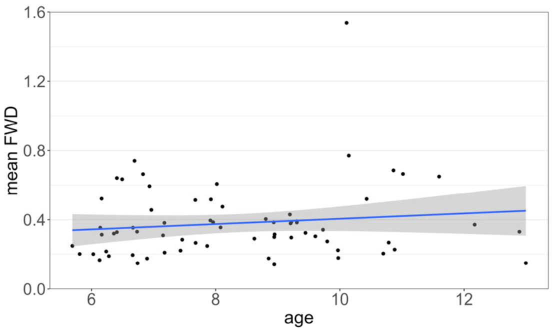


*Note.* This figure shows the mean framewise displacement value for each subject depending on age. In blue is the correlation line including the error shaded in grey.

#### A.5 Modelling: simulation and recovery

Based on the recommendations by Wilson & Collins (2019), we simulated data and recovered the parameters from the simulated data before fitting the real data. For this, we created 100 data sets with 44 trials based on our trial structures, where one of the 16 possibilities was chosen randomly. Then, for each dataset, we randomly chose values for each free parameter from a uniform distribution $U \sim uniform(0,1)$. The learning rate $\eta$ ranged from 0 to 1, the non-decision time $\tau$ from 0.3 to 3.0, the drift weight $v_{mod}$ from 0 to 15, and the decision boundary $a$ from 1 to 5. With the random parameters for each dataset, artificial response and reaction time data was calculated. This artificial data was then given into the fitting algorithm to determine whether we can recover the randomly chosen parameters for each data set. For the parameter estimation, the same upper and lower boundaries were administered to each parameter, except the non-decision time, where the upper boundary was limited to the shortest reaction time (> 200ms) per run. Parameters were recovered for each data set of 44 trials separately. The results of the recovery can be seen in **Figure A5A** and show that we can successfully recover all four free parameters with correlations between the simulated and recovered parameter ranging from *R* = .69 to *R* = 1.0 (*p*s < .001). Additionally, **Figure A5B** illustrates that the correlations within each parameter are high but around zero between parameters. Since all parameters could be recovered in the full range of the simulated values, the same interval was also used for parameter fitting on the real data.

**Figure A5**

*Comparison of simulated vs fitted parameters*

**A Scatter plot for simulated and recovered parameter values** **B Correlation plot for simulated and recovered parameter values**

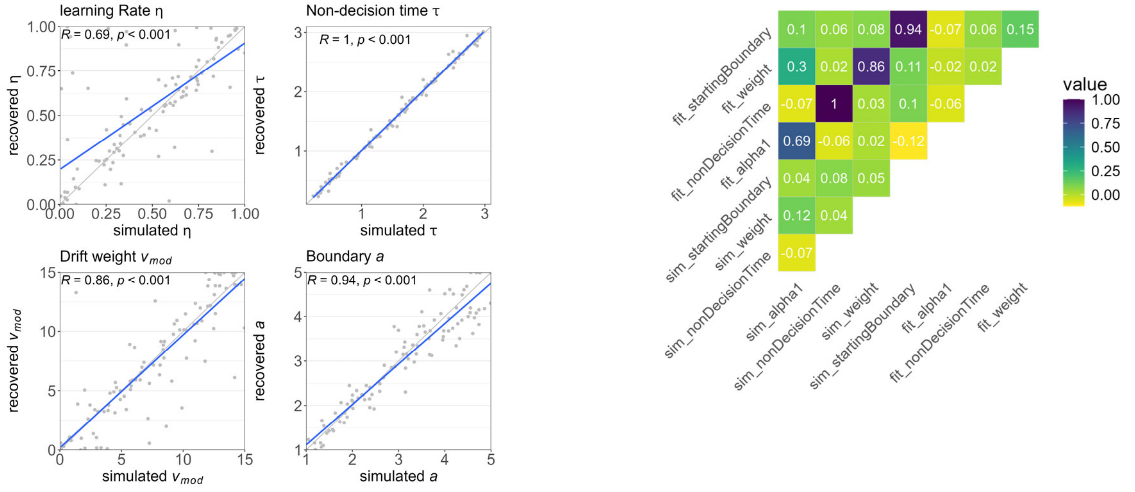


*Note.* Panel **A** shows the simulated vs fitted parameter values for learning rate, non-decision time, drift weight, and boundary separation (top left to top right). Panel **B** shows the correlations between simulated and fitted parameter values for each combination. There are high correlations within corresponding parameters (e.g. $R$ = 1.0 for simulated and fitted non-decision time) and low correlations between different parameters (e.g. $R$ = -.02 between simulated learning rate and fitted non-decision time). Higher correlations are depicted in darker colours.

#### A.6 Preprocessing of behavioural data

Following the exclusion of nine runs with over 20% omissions, a total of 7’920 trials were available for statistical analysis. Additionally, trials with no reaction (3.19%), a reaction time (RT) faster than 200 ms (0.10%), or slower than three times the standard deviation within each run (0.86%) were excluded from any further analyses, as recommended by Berger & Kiefer (2021), leading to 7’594 valid trials.

#### A.7 Processing of the ROI masks

The terms “multisensory”, “value”, and “prediction error” were each entered in the term based meta-analysis tab on neurosynth (www.neurosynth.org) and then the uniformity test activation maps were downloaded with the upper threshold set to 8, 4, or 3 for the “value”, “prediction error”, and “multisensory” map, respectively. The uniformity maps showed a higher overlap with the brain regions summarised in the literature compared to the association test map. Additionally, the association map for the term “tactile” with an upper threshold of 4 was downloaded. The maps were processed to create masks for ROI analyses. First, each map was resampled to match the dimensions of the functional scans. Then binary maps were created from each mask were all voxels exceeding a certain z-values were set to 1 and the rest of the voxels set to 0. The threshold for the “value” map was ≥ 7, for “prediction error” mask ≥ 5.5, and for the “multisensory” and “tactile” maps ≥ 4. This step ensured that distinct subclusters were identified in each hemisphere without blending multiple regions. Next, MarsBaR (version 0.45) was used to create a binary mask for each subcluster of each binary network mask. Only subclusters with at least 25 voxels were selected for further analysis. These selected subclusters were then combined into a numbered, labelled ROI file using MarsBaR. To create a mask for multisensory integration that accounted for both audio-visual and tactile-visual regions, the subclusters from the “multisensory” and “tactile” masks (each with at least 25 voxels) were combined into a single numbered ROI. Finally, each subcluster within the masks was appropriately labelled to facilitate further analyses.

### B Supplementary Material: Results

#### B.1 Analyses without diagnosed subjects

The analyses of the key findings were also done with the subgroup of subjects without a clinical diagnosis (ADHD, DD, DLD) and/or a clinically significant T-value on the CBCL/6-18R, see Supplementary Material A.1). We excluded 10 subjects ($M$ = 9.12 ± 1.54 years, 3 girls), which lead to a sample of 57 children ($M$ = 8.22 ± 1.85 years, 33 girls).

*Drift Rate:* The effects on the drift rate did not differ from the effects in the whole group.

*Age Effects during Stimulus and Feedback Processing:* During AV and TV stimulus processing, brain activation correlated with age is largely comparable to the results in the whole group. The positive main effect of age during stimulus processing is also comparable. The main effect of age and the interaction effects between age and modality on stimulus processing were identical, with an additional negative main effect of age in the subgroup in the cerebellum. Brain activation positively correlated with age during AV feedback processing was largely comparable, but without the cluster in the left middle frontal gyrus. The negative correlation with age during AV feedback processing was comparable to the whole group with an additional cluster in the right supramarginal gyrus. During TV feedback processing, the positive correlation with age was comparable but exhibited an additional cluster in the left superior medial gyrus in the subgroup. The main effect of age during feedback processing showed a cluster in the left Rolandic operculum and an additional negative main effect of age in the inferior temporal gyrus. The interaction effect during FB processing with a stronger age effect in AV compared to TV runs was comparable in the subgroup without the cluster in the angular gyrus. In the subgroup, we observed a stronger age effect in TV compared to AV runs in the right supramarginal gyrus which we did not observe in the whole group (see **Figure B1** the **Table B1.1**).

*Modulation of BOLD Signal by RPE:* Brain activation during RPE processing is comparable between the two groups. The correlation between age and RPE processing in the middle frontal gyrus did not survive cluster thresholding.

*RPE Network ROI Analyses:* The results in the RPE network ROI showed the same main effect as in the whole group. The interaction effect between age and modality in the subgroup only showed a trend (see **Table B1.2**).

**Figure B1**

*Age-effects on brain activation during stimulus and feedback processing*

**A Main effect of age for stimulus and feedback processing**

Stimulus processing | Feedback processing

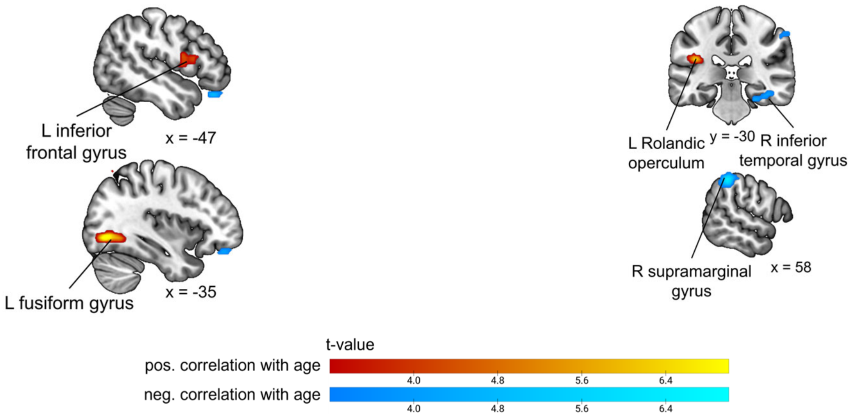


**B Interaction effect of age and modality for stimulus and feedback processing**

Stimulus processing | Feedback processing

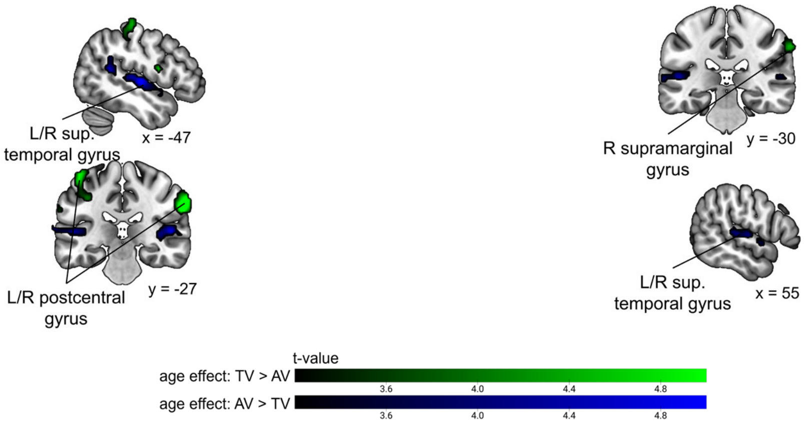


*Note.* Panel **A** shows the positive (in warm colours) and negative (cold colours) main effect of age during stimulus (left column) and feedback processing (right column) for AV and TV runs. And panel **B** shows the interaction effect of age and modality for stimulus (left) and feedback (right) processing, showing a stronger age effect in AV runs in blue, and stronger age effect in TV runs in green. Cluster defining threshold $p$ < .001 (unc.), $p_{FWEc}$ = .05, family-wise error corrected. AV = audio-visual, TV = tactile-visual, L = left, R = right, sup. = superior, neg. = negative, pos. = positive.

**Table B1.1**

*Significant clusters showing higher or lower activation with higher age during AV and TV stimulus processing*

| Contrast | Brain Area | MNI Coordinates | | | Cluster size | T-value | Peak-level $p_{unc}$ | Cluster-level $p_{FWE}$ |
|---|---|---|---|---|---|---|---|---|
| | | x | y | z | | | | |
| **Stimulus Processing** | | | | | | | | |
| **Main effect of age** | L Fusiform Gyrus | -35 | -72 | -6 | 145 | 8.25 | <.001 | .003 |
| | L Posterior-Medial Frontal | 1 | 6 | 69 | 173 | 5.77 | <.001 | .001 |
| | L IFG (p. Opercularis) | -50 | 6 | 15 | 82 | 5.06 | <.001 | .034 |
| | L Middle Occipital Gyrus | -23 | -72 | 42 | 148 | 4.85 | <.001 | .002 |
| | L Cerebellum (VI) | -17 | -78 | -18 | 131 | -5.47 | <.001 | .004 |
| | Location not in atlas | -47 | 33 | -24 | 75 | -4.84 | <.001 | .047 |
| **Interaction effect age: AV > TV** | R Superior Temporal Gyrus | 58 | -18 | 3 | 439 | 5.90 | <.001 | <.001 |
| | L Superior Temporal Gyrus | -44 | -18 | 3 | 220 | 5.33 | <.001 | <.001 |
| | L Middle Temporal Gyrus | -47 | -48 | 15 | 74 | 4.53 | <.001 | .049 |
| | R SupraMarginal Gyrus | 64 | -27 | 33 | 240 | -6.14 | <.001 | <.001 |
| | L IFG (p. Opercularis) | -53 | 3 | 15 | 106 | -6.07 | <.001 | .012 |
| | L Postcentral Gyrus | -44 | -24 | 63 | 250 | -5.46 | <.001 | <.001 |
| | L Postcentral Gyrus | -62 | -21 | 33 | 97 | -4.64 | <.001 | .018 |
| **Feedback Processing** | | | | | | | | |
| **Main effect of age** | L Rolandic Operculum | -38 | -30 | 21 | 118 | 6.56 | <.001 | .007 |
| | R SupraMarginal Gyrus | 58 | -39 | 45 | 128 | -6.56 | <.001 | .005 |
| | R Inferior Temporal Gyrus | 46 | -36 | -15 | 107 | -5.23 | <.001 | .011 |
| **Interaction effect age: AV > TV** | L Superior Temporal Gyrus | -41 | -24 | 9 | 222 | 4.63 | <.001 | <.001 |
| | R Superior Temporal Gyrus | 43 | -24 | 6 | 164 | 4.32 | <.001 | .001 |
| **TV > AV** | R SupraMarginal Gyrus | 67 | -30 | 39 | 77 | -4.49 | <.001 | .042 |

Note. Only the main peak is shown in this table. Regions were automatically labelled using the AnatomyToolbox atlas. x, y, and z = Montreal Neurological Institute (MNI) coordinates in the left-right, anterior-posterior, and inferior-superior dimensions, respectively. Cluster defining threshold $p$ < .001 (unc.), $p_{FWEc}$ = .05, family-wise error corrected. L = left, R = right, AV = audio-visual, TV = tactile-visual, IFG = inferior frontal gyrus.

**Table B1.2**

*Adjusted p-values for the effects of modality, age, and their interaction in the RPE network ROI*

| Model | Term | NumDF | DenDF | statistic | *p* unc. | *p* | $\eta^2_p$ | 95% CI |
|---|---|---|---|---|---|---|---|---|
| Whole ROI | **modality** | **1** | **55.00** | **8.26** | **.006** | **.023*** | **.13** | **[.02, 1.00]** |
| | **age** | **1** | **55.00** | **11.15** | **.002** | **.007**** | **.17** | **[.04, 1.00]** |
| | modality × age | 1 | 55.00 | 5.71 | .020 | .061. | .09 | [.01, 1.00] |
| L Anterior Insula | modality | 1 | 55.00 | 2.96 | .091 | .150 | .05 | [.00, 1.00] |
| | **age** | **1** | **55.00** | **15.54** | **.000** | **.003**** | **.22** | **[.08, 1.00]** |
| | modality × age | 1 | 55.00 | 4.11 | .048 | .114 | .07 | [.00, 1.00] |
| R Anterior Insula | modality | 1 | 55.00 | 2.91 | .094 | .150 | .05 | [.00, 1.00] |
| | **age** | **1** | **55.00** | **30.47** | **.000** | **< .001***** | **.36** | **[.19, 1.00]** |
| | modality × age | 1 | 55.00 | 1.37 | .246 | .269 | .02 | [.00, 1.00] |
| L Precentral Gyrus | modality | 1 | 55.00 | 4.25 | .044 | .114 | .07 | [.00, 1.00] |
| | age | 1 | 55.00 | 3.82 | .056 | .118 | .06 | [.00, 1.00] |
| | modality × age | 1 | 55.00 | 1.54 | .219 | .263 | .03 | [.00, 1.00] |
| R Precentral Gyrus | modality | 1 | 55.00 | 2.11 | .152 | .202 | .04 | [.00, 1.00] |
| | **age** | **1** | **55.00** | **12.65** | **.001** | **.005**** | **.19** | **[.06, 1.00]** |
| | modality × age | 1 | 55.00 | 2.59 | .114 | .160 | .04 | [.00, 1.00] |
| L/R suppl. motor | modality | 1 | 55.00 | 2.01 | .162 | .205 | .04 | [.00, 1.00] |
| Cortex | **age** | **1** | **55.00** | **14.08** | **.000** | **.003**** | **.20** | **[.07, 1.00]** |
| | modality × age | 1 | 55.00 | 3.16 | .081 | .150 | .05 | [.00, 1.00] |
| L Ventral Striatum | modality | 1 | 55.00 | 2.65 | .109 | .160 | .05 | [.00, 1.00] |
| | age | 1 | 55.00 | 1.42 | .238 | .269 | .03 | [.00, 1.00] |
| | modality × age | 1 | 55.00 | 0.56 | .459 | .459 | .01 | [.00, 1.00] |
| R Ventral Striatum | **modality** | **1** | **55.00** | **7.57** | **.008** | **.028*** | **.12** | **[.02, 1.00]** |
| | age | 1 | 55.00 | 0.90 | .348 | .363 | .02 | [.00, 1.00] |
| | modality × age | 1 | 55.00 | 3.72 | .059 | .118 | .06 | [.00, 1.00] |

*Note*. Type III Analysis of Variance table with Satterthwaite's method. Highlighted in grey are the significant effects after correction for multiple comparisons using the FDR method. L = left, R = right, RPE = reward prediction error, ROI = region of interest, suppl. = supplementary, NumDF = numerator degrees of freedom, DenDF = denominator degrees of freedom, *p* unc. = uncorrected *p*-value, $\eta^2_p$ = partial eta-squared effect sizes, CI = confidence interval. Signif. codes: 0 '***' 0.001 '**' 0.01 '*' 0.05 '.' 0.1 ' ' 1. Interpretation of effect sizes: $\eta^2_P < .01$ very small, $.01 \leq \eta^2_P < .06$ small, $.06 \leq \eta^2_P < .14$ medium, $\eta^2_P \geq .14$ large (Field, 2013).

#### B.2 Behavioural results

As an overview of general task performance, **Table B2.1** shows mean values of reaction times for correct trials, mean number of omissions and outliers in reaction time, and median accuracies and inter-quartile range (IQR) per modality.

**Table B2.1**

Task performance

| | **AV** | **TV** |
|---|---|---|
| **Accuracy** [%; Md (IQR)] | 65.91 (24.70) | 65.12 (23.58) |
| **Reaction Time** correct trials [s] | 2.26 (0.37) | 2.49 (0.35) |
| **Omissions** [%] | 2.85 (3.87) | 4.31 (4.60) |
| **Outliers** [%] | 0.80 (1.22) | 1.10 (1.24) |

Note. The following measures of task performance are provided. Mean and standard deviation values are given unless otherwise indicated. Accuracy = number of correct trials divided by total trials not counting omissions. Omissions = number of non-responses expressed as a percentage per run. Outliers = combined amount of fast (< 200ms) and slow (> mean + 3*standard deviation) reaction times, expressed as a percentage per run. Md = median, IQR = inter-quartile range, AV = audio-visual, TV = tactile-visual.

**Table B2.2**

*Post-hoc tests of the effect of bin on reaction time and accuracy*

| Model | contrast | estimate | SE | *df* | *t* | *p* |
|---|---|---|---|---|---|---|
| reaction time ~ | 1st - 2nd | 0.04 | 0.03 | 464.00 | 1.07 | .708 |
| bin | **1st - 3rd** | **0.15** | **0.03** | **464.00** | **4.36** | **< .001***** |
| | **1st - 4th** | **0.13** | **0.03** | **464.00** | **3.78** | **.001**** |
| | **2nd - 3rd** | **0.11** | **0.03** | **464.00** | **3.29** | **.006**** |
| | **2nd - 4th** | **0.09** | **0.03** | **464.00** | **2.71** | **.035*** |
| | 3rd - 4th | -0.02 | 0.03 | 464.00 | -0.58 | .939 |
| accuracy ~ | 1st - 2nd | -0.02 | 0.02 | 464.00 | -1.13 | .671 |
| bin | **1st - 3rd** | **-0.09** | **0.02** | **464.00** | **-4.92** | **< .001***** |
| | **1st - 4th** | **-0.10** | **0.02** | **464.00** | **-5.18** | **< .001***** |
| | **2nd - 3rd** | **-0.07** | **0.02** | **464.00** | **-3.79** | **.001***** |
| | **2nd - 4th** | **-0.08** | **0.02** | **464.00** | **-4.05** | **< .001***** |
| | 3rd - 4th | -0.00 | 0.02 | 464.00 | -0.25 | .994 |

*Note*. Post-hoc pairwise comparisons for the effect of the specified factor on the dependent variable. Estimates represent the mean difference between conditions. SE = standard error, *df* = degrees of freedom, *p* values are Bonferroni-corrected for multiple comparisons.

#### 1 B.3 Whole-brain stimulus processing

**Table B3.1**

*Clusters with significant correlation between activation during AV and TV stimulus processing and age*

| Contrast | Brain Area | MNI Coordinates | | | Cluster size | T-value | Peak-level $p_{unc}$ | Cluster-level $p_{FWE}$ |
|---|---|---|---|---|---|---|---|---|
| | | x | y | z | | | | |
| **AV** | L IFG (p. Opercularis) | -56 | 21 | 39 | 541 | 7.09 | <.001 | <.001 |
| | L Inferior Occipital Gyrus | -35 | -78 | -3 | 1821 | 6.97 | <.001 | <.001 |
| | L Middle Cingulate Cortex | -5 | 9 | 48 | 279 | 6.62 | <.001 | <.001 |
| | R Insula Lobe | 34 | 21 | 6 | 86 | 5.19 | <.001 | .032 |
| | R Temporal Pole | 67 | 6 | 0 | 293 | 4.94 | <.001 | <.001 |
| | L Superior Temporal Gyrus | -41 | -15 | -3 | 196 | 4.48 | <.001 | .001 |
| | R Cerebellum (Crus 2) | 7 | -84 | -30 | 87 | 4.39 | <.001 | .031 |
| | R Postcentral Gyrus | 25 | -48 | 72 | 337 | -6.12 | <.001 | <.001 |
| | L Precuneus | -14 | -54 | 69 | 305 | -6.09 | <.001 | <.001 |
| | R SupraMarginal Gyrus | 67 | -27 | 33 | 310 | -5.73 | <.001 | <.001 |
| | L Calcarine Gyrus | -14 | -57 | 15 | 115 | -5.41 | <.001 | .010 |
| | R ParaHippocampal Gyrus | 19 | -15 | -21 | 104 | -5.31 | <.001 | .015 |
| | L Middle Temporal Gyrus | -53 | -9 | -15 | 145 | -5.23 | <.001 | .003 |
| | R Superior Medial Gyrus | 13 | 60 | 18 | 624 | -5.10 | <.001 | <.001 |
| | L ParaHippocampal Gyrus | -20 | -15 | -21 | 95 | -4.77 | <.001 | .022 |
| | L Rolandic Operculum | -59 | -3 | 18 | 111 | -4.38 | <.001 | .012 |
| | R IFG (p. Orbitalis) | 28 | 30 | -15 | 89 | -4.28 | <.001 | .029 |
| **TV** | L IFG (p. Triangularis) | -44 | 27 | 30 | 756 | 9.57 | <.001 | <.001 |
| | L Angular Gyrus | -32 | -51 | 42 | 2628 | 8.46 | <.001 | <.001 |
| | R Middle Frontal Gyrus | 37 | 30 | 24 | 621 | 6.24 | <.001 | <.001 |
| | L Inferior Occipital Gyrus | -41 | -66 | -6 | 690 | 6.16 | <.001 | <.001 |
| | R Inferior Occipital Gyrus | 37 | -90 | 0 | 144 | 4.78 | <.001 | .002 |
| | Location not in atlas | -5 | -27 | 27 | 76 | 4.77 | <.001 | .042 |

| | | | | | | | | |
|---|---|---|---|---|---|---|---|---|
| | R Middle Cingulate Cortex | 16 | -33 | 45 | 1040 | -7.04 | <.001 | <.001 |
| | R Medial Temporal Pole | 52 | 6 | -27 | 448 | -5.66 | <.001 | <.001 |
| | L Middle Temporal Gyrus | -56 | 0 | -18 | 401 | -5.63 | <.001 | <.001 |
| | R Middle Temporal Gyrus | 46 | -72 | 24 | 111 | -5.39 | <.001 | .009 |
| | L SupraMarginal Gyrus | -65 | -42 | 36 | 155 | -5.38 | <.001 | .002 |
| | L Calcarine Gyrus | -11 | -54 | 12 | 132 | -5.16 | <.001 | .004 |
| | L ParaHippocampal Gyrus | -29 | -30 | -12 | 129 | -5.05 | <.001 | .004 |
| | L Middle Temporal Gyrus | -62 | -27 | 0 | 74 | -5.00 | <.001 | .046 |
| | R Fusiform Gyrus | 22 | -39 | -12 | 124 | -4.80 | <.001 | .005 |
| **Main Effect of age** | L Fusiform Gyrus | -35 | -72 | -6 | 278 | 8.09 | <.001 | <.001 |
| | L IFG (p. Opercularis) | -47 | 6 | 6 | 192 | 6.38 | <.001 | <.001 |
| | L Middle Occipital Gyrus | -23 | -69 | 39 | 95 | 5.65 | <.001 | .020 |
| | R Posterior-Medial Frontal | 4 | 6 | 54 | 186 | 5.16 | <.001 | .001 |
| **Interaction Effect age: AV > TV** | R Superior Temporal Gyrus | 58 | -18 | 3 | 620 | 5.75 | <.001 | <.001 |
| | L Superior Temporal Gyrus | -47 | -15 | 3 | 350 | 5.17 | <.001 | <.001 |
| **TV > AV** | R SupraMarginal Gyrus | 64 | -27 | 30 | 219 | -5.75 | <.001 | <.001 |
| | L IFG (p. Opercularis) | -53 | 3 | 15 | 85 | -5.16 | <.001 | .031 |
| | L Postcentral Gyrus | -44 | -24 | 63 | 266 | -5.14 | <.001 | <.001 |
| | L Postcentral Gyrus | -62 | -21 | 33 | 88 | -4.55 | <.001 | .027 |

*Note*. Only the main peak is shown in this table. Regions were automatically labelled using the AnatomyToolbox atlas. x, y, and z = Montreal Neurological Institute (MNI) coordinates in the left-right, anterior-posterior, and inferior-superior dimensions, respectively. Cluster defining threshold $p$ < .001 (unc.), $p_{FWEc}$ = .05, family-wise error corrected. L = left, R = right, AV = audio-visual, TV = tactile-visual, IFG = inferior frontal gyrus.

During AV stimulus presentation, auditory, visual, and fronto-central regions show significant activation. During TV stimulus processing, mainly, a left and right somatosensory cluster extending into frontal regions and a visual cluster is observed. All clusters can be seen in **Table B3.2** and **Figure B3**.

**Figure B3**

Whole-brain stimulus processing during AV and TV runs

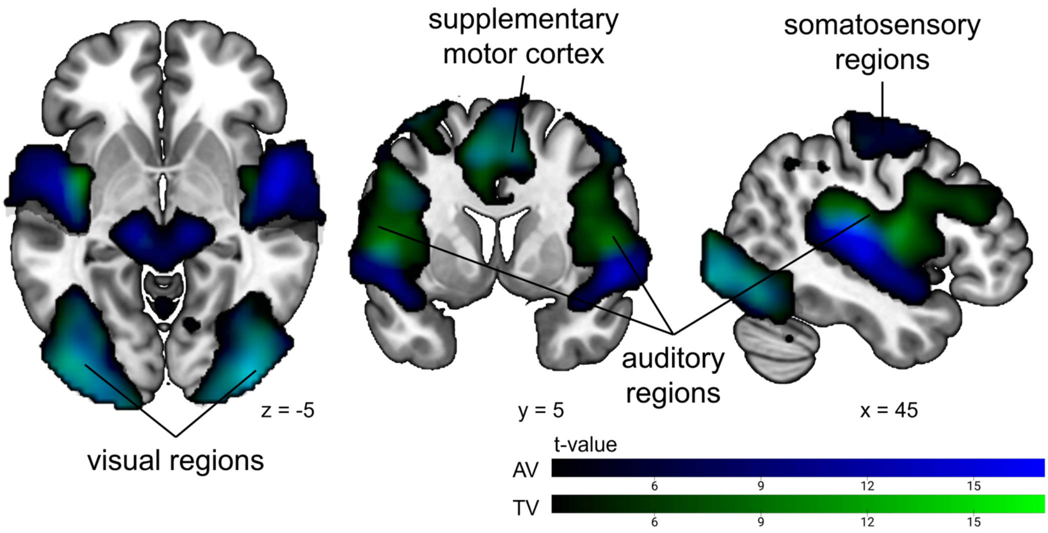


*Note.* Stimulus processing during AV runs is shown in blue, during TV runs in green. Overlapping regions are coloured in light blue. Cluster defining threshold $p$ < .001 (unc.), $p_{FWEc}$ = .05, family-wise error corrected. AV = audio-visual, TV = tactile-visual.

**Table B3.2**

*Clusters with significant activation during AV and TV stimulus processing*

| Contrast | Brain Area | MNI Coordinates | | | Cluster size | T-value | Peak-level | Cluster-level |
|---|---|---|---|---|---|---|---|---|
| | | x | y | z | | | $p_{unc}$ | $p_{FWE}$ |
| **AV** | R Superior Temporal Gyrus | 64 | -27 | 12 | 2066 | 18.53 | <.001 | <.001 |
| | L Superior Temporal Gyrus | -47 | -21 | 9 | 7968 | 17.16 | <.001 | <.001 |
| | L Middle Cingulate Cortex | -8 | 9 | 48 | 838 | 12.16 | <.001 | <.001 |
| | L Thalamus | -11 | -27 | -6 | 622 | 11.62 | <.001 | <.001 |
| | L Insula Lobe | -29 | 27 | 6 | 873 | 8.68 | <.001 | <.001 |
| | R Insula Lobe | 31 | 24 | 3 | 79 | 6.56 | <.001 | .044 |
| | R Middle Frontal Gyrus | 49 | 0 | 57 | 553 | 6.50 | <.001 | <.001 |
| **TV** | L Insula Lobe | -41 | -6 | 6 | 5741 | 16.38 | <.001 | <.001 |
| | R Inferior Occipital Gyrus | 40 | -90 | -6 | 1803 | 13.93 | <.001 | <.001 |
| | L Linual Gyrus | -35 | -90 | - 12 | 1768 | 13.54 | <.001 | <.001 |
| | R Rolandic Operculum | 49 | -21 | 21 | 2508 | 12.27 | <.001 | <.001 |
| | R Angular Gyrus | 28 | -60 | 42 | 472 | 9.39 | <.001 | <.001 |
| | L Thalamus | -14 | -27 | 9 | 185 | 7.40 | <.001 | .001 |
| | Location not in atlas | 4 | -30 | 27 | 114 | 5.99 | <.001 | .008 |
| | R Precentral Gyrus | 52 | -18 | 51 | 118 | 4.68 | <.001 | .007 |

*Note*. Only the main peak is shown in this table. Regions were automatically labelled using the AnatomyToolbox atlas. x, y, and z = Montreal Neurological Institute (MNI) coordinates in the left-right, anterior-posterior, and inferior-superior dimensions, respectively. Cluster defining threshold $p$ < .001 (unc.), $p_{FWEc}$ = .05, family-wise error corrected. L = left, R = right, AV = audio-visual, TV = tactile-visual.

#### B.4 Whole-brain feedback processing

A main effect of age during feedback processing was observed exclusively in the left cerebellum (lobule VI), with older children showing higher activation across sensory modalities. Activation of the cerebellum has been reported previously during performance feedback processing, error processing (Peterburs et al., 2015, 2018), reversal learning (Peterburs et al., 2018; Waltz et al., 2013) as well as for (verbal) working memory processes (Stoodley et al., 2012). These findings have to be interpreted with caution, since we did not have cerebellum coverage for all participants.

**Table B4.1**

*Clusters with significant correlation between activation during AV and TV feedback processing and age*

| Contrast | Brain Area | MNI Coordinates | | | Cluster size | T-value | Peak-level $p_{unc}$ | Cluster-level $p_{FWE}$ |
|---|---|---|---|---|---|---|---|---|
| | | x | y | z | | | | |
| **AV** | L Fusiform Gyrus | -32 | -75 | -6 | 397 | 5.97 | <.001 | <.001 |
| | Location not in atlas | 73 | -39 | 6 | 457 | 5.67 | <.001 | <.001 |
| | R Inferior Occipital Gyrus | 37 | -93 | 3 | 213 | 5.48 | <.001 | <.001 |
| | L Middle Frontal Gyrus | -41 | 12 | 54 | 119 | 5.15 | <.001 | .009 |
| | L Heschls Gyrus | -47 | -21 | 12 | 461 | 5.04 | <.001 | <.001 |
| | R Pallidum | 22 | -3 | 0 | 156 | -4.94 | <.001 | .002 |
| | L Postcentral Gyrus | -32 | -42 | 60 | 117 | -4.91 | <.001 | .009 |
| | R Rolandic Operculum | 49 | 3 | 15 | 84 | -4.91 | <.001 | .036 |
| **TV** | L Inferior Occipital Gyrus | -32 | -75 | -3 | 243 | 5.55 | <.001 | <.001 |
| | L Superior Frontal Gyrus | -20 | 57 | 33 | 93 | 4.83 | <.001 | .021 |
| **Neg. main effect of age** | R SupraMarginal Gyrus | 58 | -39 | 45 | 128 | -6.26 | <.001 | .005 |
| **Interaction effect age: AV > TV** | L Superior Temporal Gyrus | -44 | -24 | 9 | 331 | 5.03 | <.001 | <.001 |
| | R Angular Gyrus | 55 | -63 | 30 | 158 | 4.79 | <.001 | .002 |
| | R Temporal Pole | 52 | 0 | -3 | 251 | 4.58 | <.001 | <.001 |

Note. Only the main peak is shown in this table. Regions were automatically labelled using the AnatomyToolbox atlas. x, y, and z = Montreal Neurological Institute (MNI) coordinates in the left-right, anterior-posterior, and inferior-superior dimensions, respectively. Cluster defining threshold $p$ < .001 (unc.), $p_{FWEc}$ = .05, family-wise error corrected. L = left, R = right, AV = audio-visual, TV = tactile-visual, neg. = negative.

When looking at feedback processing during AV and TV runs combined, mainly visual regions showed increased brain activation. During AV feedback processing, additional clusters in auditory regions show higher activation. There is no additional cluster in TV runs (more details in **Figure B4** and **Table B4.2**).

**Figure B4**

*Whole-brain feedback processing during AV, TV runs, and AV + TV runs combined*

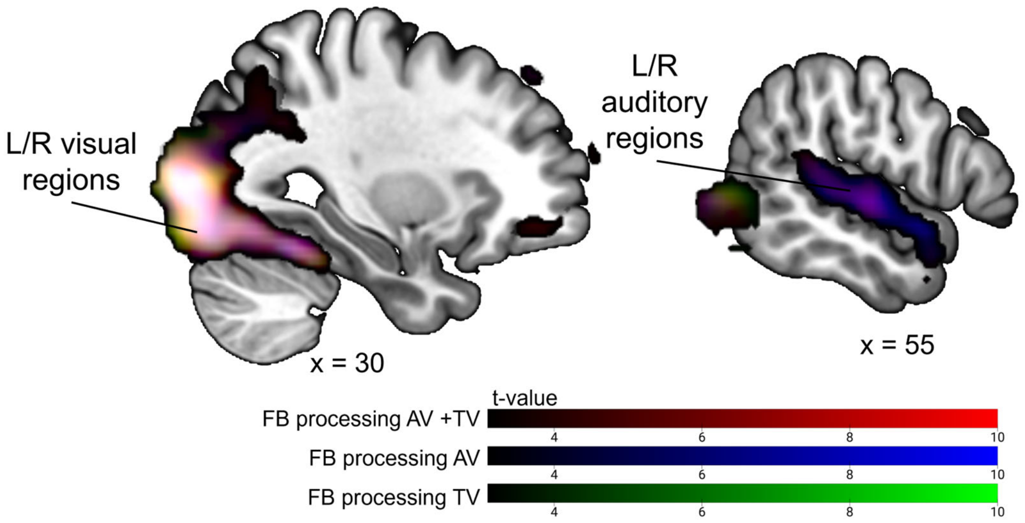


*Note.* Feedback processing during AV and TV runs combined in red, AV runs in blue, and TV runs in green. Cluster defining threshold $p$ < .001 (unc.), $p_{FWEc}$ = .05, family-wise error corrected. AV = audio-visual, TV = tactile-visual, L = left, R = right, FB = feedback

**Table B4.2**

*Significant clusters during feedback processing*

| Contrast | Brain Area | MNI Coordinates | | | Cluster size | T-value | Peak-level $p_{unc}$ | Cluster-level $p_{FWE}$ |
|---|---|---|---|---|---|---|---|---|
| | | x | y | z | | | | |
| **AV + TV** | R Inferior Occipital Gyrus | 37 | -84 | -9 | 2795 | 15.98 | <.001 | <.001 |
| | L Fusiform Gyrus | -32 | -84 | -9 | 2946 | 14.95 | <.001 | <.001 |
| | R Superior Medial Gyrus | 10 | 48 | 54 | 735 | 6.45 | <.001 | <.001 |
| | L Brainstem | -8 | -30 | -6 | 116 | 6.11 | <.001 | .009 |
| | R Middle Orbital Gyrus | 22 | 33 | -12 | 87 | 4.82 | <.001 | .030 |
| **AV** | L Cerebellum (VI) | -38 | -72 | -15 | 1658 | 13.38 | <.001 | <.001 |
| | R Middle Occipital Gyrus | 28 | -90 | 6 | 1907 | 13.19 | <.001 | <.001 |
| | R Superior Temporal Gyrus | 61 | -18 | 0 | 712 | 8.32 | <.001 | <.001 |
| | L Middle Temporal Gyrus | -62 | -39 | 9 | 970 | 8.23 | <.001 | <.001 |
| | L Brainstem | -5 | -33 | -9 | 179 | 7.08 | <.001 | .001 |
| | R Superior Medial Gyrus | 7 | 48 | 54 | 359 | 5.68 | <.001 | .014 |
| **TV** | R Inferior Occipital Gyrus | 40 | -78 | -6 | 1923 | 11.73 | <.001 | <.001 |
| | L Fusiform Gyrus | -35 | -84 | -12 | 1564 | 11.15 | <.001 | <.001 |
| | L Rolandic Operculum | -38 | -39 | 21 | 560 | 6.53 | <.001 | <.001 |
| | R Superior Medial Gyrus | 13 | 63 | 33 | 264 | 5.80 | <.001 | <.001 |

Note. Only the main peak is shown in this table. Regions were automatically labelled using the AnatomyToolbox atlas. x, y, and z = Montreal Neurological Institute (MNI) coordinates in the left-right, anterior-posterior, and inferior-superior dimensions, respectively. Cluster defining threshold $p$ < .001 (unc.), $p_{FWEc}$ = .05, family-wise error corrected. L = left, R = right, AV = audio-visual, TV = tactile-visual.

#### B.5 Reward prediction error processing during AV and TV runs

RPEs positively modulated brain activation in the ventral striatum, precuneus, and frontal regions during AV runs. In contrast, inferior frontal and insular regions showed negative modulation by RPEs in the same condition. During TV runs, RPEs positively modulated activation in the ventral striatum, while insular regions exhibited negative modulation (for more details see **Table B5**).

**Table B5**

*Clusters with brain activation during feedback processing modulated by RPEs*

| Contrast | Brain Area | MNI Coordinates | | | Cluster size | T-value | Peak-level $p_{unc}$ | Cluster-level $p_{FWE}$ |
|---|---|---|---|---|---|---|---|---|
| | | x | y | z | | | | |
| **AV modulated by RPE** | R Calcarine Gyrus | 7 | -57 | 15 | 989 | 7.32 | <.001 | <.001 |
| | L Angular Gyrus | -44 | -75 | 33 | 210 | 6.18 | <.001 | <.001 |
| | R ParaHippocampal Gyrus | 22 | -21 | -15 | 483 | 5.74 | <.001 | <.001 |
| | Location not in atlas | 25 | -21 | 27 | 95 | 5.35 | <.001 | .022 |
| | L Middle Frontal Gyrus | -23 | 6 | 57 | 181 | 5.03 | <.001 | .001 |
| | R Rectal Gyrus | 7 | 42 | -12 | 135 | 4.99 | <.001 | .005 |
| | R Postcentral Gyrus | 58 | -24 | 48 | 238 | 4.56 | <.001 | <.001 |
| | R Insula Lobe | 43 | 15 | -3 | 1838 | -10.45 | <.001 | <.001 |
| | R Middle Cingulate Cortex | 7 | 21 | 39 | 1645 | -9.43 | <.001 | <.001 |
| | L IFG (p. Orbitalis) | -32 | 27 | 0 | 1072 | -7.59 | <.001 | <.001 |
| | R Middle Temporal Gyrus | 52 | -33 | -3 | 587 | -6.69 | <.001 | <.001 |
| | R Cerebellum (VI) | 22 | -69 | -12 | 218 | -5.28 | <.001 | <.001 |
| | L Cerebellum (Crus 1) | -44 | -57 | -30 | 272 | -5.06 | <.001 | <.001 |
| | L Superior Temporal Gyrus | -56 | -48 | 21 | 96 | -4.77 | <.001 | .021 |
| **TV modulated by RPE** | L Olfactory cortex | -11 | 9 | -9 | 919 | 7.86 | <.001 | <.001 |
| | L Superior Frontal Gyrus | -20 | 33 | 45 | 141 | 6.09 | <.001 | .003 |
| | Location not in atlas | -23 | -54 | 21 | 559 | 5.10 | <.001 | <.001 |
| | Location not in atlas | 19 | 45 | 0 | 194 | 4.86 | <.001 | <.001 |
| | L Cerebellum (Crus 1) | -35 | -54 | -33 | 283 | -8.04 | <.001 | <.001 |
| | R Posterior-Medial Frontal | 10 | 18 | 60 | 1051 | -6.50 | <.001 | <.001 |
| | R Cerebellum (Crus 1) | 37 | -57 | -30 | 509 | -6.32 | <.001 | <.001 |
| | R IFG (p. Orbitalis) | 40 | 30 | 0 | 1034 | -6.24 | <.001 | <.001 |
| | L IFG (p. Orbitalis) | -32 | 18 | -6 | 431 | -6.12 | <.001 | <.001 |
| | L Middle Frontal Gyrus | -26 | 51 | 24 | 138 | -5.99 | <.001 | .003 |
| | L Superior Temporal Gyrus | -62 | -48 | 24 | 116 | -5.49 | <.001 | .008 |
| | R Superior Frontal Gyrus | 25 | 57 | 24 | 92 | -4.44 | <.001 | .022 |

**Table B5**

*Clusters with brain activation during feedback processing modulated by RPEs*

| Contrast | Brain Area | MNI Coordinates | | | Cluster size | T-value | Peak-level $p_{unc}$ | Cluster-level $p_{FWE}$ |
|---|---|---|---|---|---|---|---|---|
| | | x | y | z | | | | |
| | R Middle Temporal Gyrus | 58 | -33 | -3 | 74 | -3.91 | <.001 | .048 |

Note. Only the main peak is shown in this table. Regions were automatically labelled using the AnatomyToolbox atlas. x, y, and z = Montreal Neurological Institute (MNI) coordinates in the left-right, anterior-posterior, and inferior-superior dimensions, respectively. Cluster defining threshold $p$ < .001 (unc.), $p_{FWEc}$ = .05, family-wise error corrected. L = left, R = right, AV = audio-visual, TV = tactile-visual, RPE = reward prediction error, IFG = inferior frontal gyrus.

#### B.6 Multisensory network ROI: effects table

**Table B6**

*Adjusted p-values for the effects of modality, age, and their interaction, and learning rate in the multisensory network ROI*

| Model | Term | NumDF | DenDF | statistic | *p* unc. | *p* | $\eta^2_p$ | 95% CI |
|---|---|---|---|---|---|---|---|---|
| whole | **modality** | **1** | **63.57** | **31.35** | **.000** | **< .001*****| **.33** | **[.18, 1.00]** |
| multisensory | **age** | **1** | **63.91** | **27.61** | **.000** | **< .001***** | **.30** | **[.15, 1.00]** |
| network | **learning rate** | **1** | **112.17** | **8.81** | **.004** | **.010**** | **.07** | **[.01, 1.00]** |
| | modality × age | 1 | 64.09 | 0.20 | .660 | .787 | .00 | [.00, 1.00] |
| L Anterior | modality | 1 | 64.64 | 0.28 | .602 | .732 | .00 | [.00, 1.00] |
| Insula | **age** | **1** | **64.82** | **10.52** | **.002** | **.006**** | **.14** | **[.03, 1.00]** |
| | learning rate | 1 | 123.29 | 1.75 | .189 | .330 | .01 | [.00, 1.00] |
| | modality × age | 1 | 65.23 | 0.44 | .512 | .666 | .01 | [.00, 1.00] |
| R Anterior | modality | 1 | 64.50 | 0.31 | .580 | .732 | .00 | [.00, 1.00] |
| Insula | **age** | **1** | **64.94** | **18.95** | **.000** | **< .001***** | **.23** | **[.09, 1.00]** |
| | learning rate | 1 | 105.55 | 0.58 | .446 | .625 | .01 | [.00, 1.00] |
| | modality × age | 1 | 64.98 | 0.29 | .594 | .732 | .00 | [.00, 1.00] |
| L Inferior | modality | 1 | 63.91 | 0.00 | .955 | .955 | .00 | [.00, 1.00] |
| Occipital | **age** | **1** | **64.38** | **20.49** | **.000** | **< .001***** | **.24** | **[.10, 1.00]** |
| Cortex | learning rate | 1 | 103.26 | 0.03 | .868 | .934 | .00 | [.00, 1.00] |
| | modality × age | 1 | 64.38 | 1.43 | .237 | .401 | .02 | [.00, 1.00] |
| R Inferior | modality | 1 | 64.61 | 2.40 | .127 | .236 | .04 | [.00, 1.00] |
| Occipital | age | 1 | 64.86 | 1.30 | .258 | .426 | .02 | [.00, 1.00] |
| Cortex | learning rate | 1 | 119.24 | 0.02 | .889 | .939 | .00 | [.00, 1.00] |
| | modality × age | 1 | 65.18 | 0.12 | .735 | .840 | .00 | [.00, 1.00] |
| L Planum | **modality** | **1** | **63.49** | **197.67** | **.000** | **< .001***** | **.76** | **[.67, 1.00]** |
| Temporale | **age** | **1** | **63.63** | **11.21** | **.001** | **.005**** | **.15** | **[.04, 1.00]** |
| | learning rate | 1 | 124.90 | 5.09 | .026 | .058. | .04 | [.00, 1.00] |
| | **modality × age** | **1** | **64.09** | **10.65** | **.002** | **.006**** | **.14** | **[.04, 1.00]** |
| R Planum | **modality** | **1** | **64.77** | **241.41** | **.000** | **< .001***** | **.79** | **[.71, 1.00]** |
| Temporale | **age** | **1** | **64.72** | **12.69** | **.001** | **.003**** | **.16** | **[.05, 1.00]** |
| | **learning rate** | **1** | **128.96** | **7.61** | **.007** | **.017*** | **.06** | **[.01, 1.00]** |
| | **modality × age** | **1** | **65.40** | **15.23** | **.000** | **.001***** | **.19** | **[.07, 1.00]** |
| L Postcentral | **modality** | **1** | **63.31** | **30.67** | **.000** | **< .001***** | **.33** | **[.18, 1.00]** |
| Gyrus | **age** | **1** | **63.55** | **19.92** | **.000** | **< .001***** | **.24** | **[.10, 1.00]** |
| | learning rate | 1 | 119.59 | 3.54 | .062 | .129 | .03 | [.00, 1.00] |
| | **modality × age** | **1** | **63.88** | **10.26** | **.002** | **.006**** | **.14** | **[.03, 1.00]** |
| R Postcentral | modality | 1 | 63.80 | 0.17 | .680 | .793 | .00 | [.00, 1.00] |
| Gyrus | **age** | **1** | **63.97** | **5.74** | **.020** | **.046*** | **.08** | **[.01, 1.00]** |
| | **learning rate** | **1** | **123.75** | **9.93** | **.002** | **.006**** | **.07** | **[.02, 1.00]** |
| | modality × age | 1 | 64.40 | 0.55 | .459 | .628 | .01 | [.00, 1.00] |
| L Precentral | modality | 1 | 64.25 | 0.86 | .357 | .555 | .01 | [.00, 1.00] |
| Gyrus | **age** | **1** | **64.75** | **58.79** | **.000** | **< .001***** | **.48** | **[.33, 1.00]** |

**Table B6**

*Adjusted p-values for the effects of modality, age, and their interaction, and learning rate in the multisensory network ROI*

| Model | Term | NumDF | DenDF | statistic | *p* unc. | *p* | $\eta^2_p$ | 95% CI |
|---|---|---|---|---|---|---|---|---|
| | learning rate | 1 | 100.15 | 0.08 | .772 | .848 | .00 | [.00, 1.00] |
| | modality × age | 1 | 64.69 | 2.11 | .151 | .273 | .03 | [.00, 1.00] |
| R Precentral | **modality** | **1** | **64.51** | **25.26** | **.000** | **< .001*** ** | **.28** | **[.14, 1.00]** |
| Gyrus | **age** | **1** | **64.54** | **16.07** | **.000** | **.001*** ** | **.20** | **[.07, 1.00]** |
| | learning rate | 1 | 128.40 | 0.62 | .432 | .621 | .00 | [.00, 1.00] |
| | **modality × age** | **1** | **65.13** | **12.78** | **.001** | **.003** ** | **.16** | **[.05, 1.00]** |
| L Superior | **modality** | **1** | **64.55** | **6.04** | **.017** | **.041* ** | **.09** | **[.01, 1.00]** |
| Parietal | **age** | **1** | **64.88** | **55.68** | **.000** | **< .001*** ** | **.46** | **[.32, 1.00]** |
| Lobe | learning rate | 1 | 113.84 | 0.10 | .754 | .844 | .00 | [.00, 1.00] |
| | modality × age | 1 | 65.09 | 2.66 | .108 | .212 | .04 | [.00, 1.00] |
| R Superior | modality | 1 | 64.43 | 4.25 | .043 | .093. | .06 | [.00, 1.00] |
| Parietal | age | 1 | 64.83 | 0.82 | .370 | .559 | .01 | [.00, 1.00] |
| Lobe | learning rate | 1 | 108.83 | 0.00 | .944 | .955 | .00 | [.00, 1.00] |
| | modality × age | 1 | 64.93 | 0.01 | .906 | .939 | .00 | [.00, 1.00] |
| Thalamus | modality | 1 | 64.55 | 1.24 | .269 | .430 | .02 | [.00, 1.00] |
| | age | 1 | 64.80 | 2.63 | .110 | .212 | .04 | [.00, 1.00] |
| | learning rate | 1 | 119.64 | 0.74 | .393 | .578 | .01 | [.00, 1.00] |
| | modality × age | 1 | 65.12 | 0.49 | .488 | .651 | .01 | [.00, 1.00] |

*Note*. Type III Analysis of Variance table with Satterthwaite's method. Highlighted in grey are the significant effects after correction for multiple comparisons using the FDR method. L = left, R = right, ROI = region of interest, NumDF = numerator degrees of freedom, DenDF = denominator degrees of freedom, *p* unc. = uncorrected *p*-value, $\eta^2_p$ = partial eta-squared effect sizes, CI = confidence interval. Signif. codes: 0 '***' 0.001 '**' 0.01 '*' 0.05 '.' 0.1 ' ' 1. Interpretation of effect sizes: $\eta^2_P < .01$ very small, $.01 \le \eta^2_P < .06$ small, $.06 \le \eta^2_P < .14$ medium, $\eta^2_P \ge .14$ large (Field, 2013).

1

#### B.7 Reward prediction error network ROI: effects table

**Table B7**

*Adjusted p-values for the effects of modality, age, and their interaction in the RPE network ROI*

| Model | Term | NumDF | DenDF | statistic | *p* unc. | *p* | $\eta^2_p$ | 95% CI |
|---|---|---|---|---|---|---|---|---|
| Whole ROI | **modality** | **1** | **65.00** | **11.21** | **.001** | **.005**** | **.15** | **[.04, 1.00]** |
| | **age** | **1** | **65.00** | **14.30** | **.000** | **.002**** | **.18** | **[.06, 1.00]** |
| | **modality × age** | **1** | **65.00** | **6.19** | **.015** | **.046*** | **.09** | **[.01, 1.00]** |
| L Anterior Insula | modality | 1 | 65.00 | 3.43 | .069 | .116 | .05 | [.00, 1.00] |
| | **age** | **1** | **65.00** | **16.99** | **.000** | **.001***** | **.21** | **[.08, 1.00]** |
| | modality × age | 1 | 65.00 | 4.86 | .031 | .068. | .07 | [.00, 1.00] |
| R Anterior Insula | modality | 1 | 65.00 | 5.26 | .025 | .060. | .07 | [.01, 1.00] |
| | **age** | **1** | **65.00** | **36.07** | **.000** | **< .001***** | **.36** | **[.21, 1.00]** |
| | modality × age | 1 | 65.00 | 2.06 | .156 | .188 | .03 | [.00, 1.00] |
| L Precentral Gyrus | modality | 1 | 65.00 | 3.72 | .058 | .107 | .05 | [.00, 1.00] |
| | age | 1 | 65.00 | 1.61 | .209 | .239 | .02 | [.00, 1.00] |
| | modality × age | 1 | 65.00 | 2.34 | .131 | .185 | .03 | [.00, 1.00] |
| R Precentral Gyrus | modality | 1 | 65.00 | 2.05 | .157 | .188 | .03 | [.00, 1.00] |
| | **age** | **1** | **65.00** | **12.27** | **.001** | **.004**** | **.16** | **[.05, 1.00]** |
| | modality × age | 1 | 65.00 | 4.12 | .047 | .093. | .06 | [.00, 1.00] |
| L/R suppl. motor | modality | 1 | 65.00 | 3.19 | .079 | .118 | .05 | [.00, 1.00] |
| Cortex | **age** | **1** | **65.00** | **18.93** | **.000** | **.001***** | **.23** | **[.09, 1.00]** |
| | modality × age | 1 | 65.00 | 5.44 | .023 | .060. | .08 | [.01, 1.00] |
| L Ventral Striatum | modality | 1 | 65.00 | 3.33 | .073 | .116 | .05 | [.00, 1.00] |
| | age | 1 | 65.00 | 2.10 | .152 | .188 | .03 | [.00, 1.00] |
| | modality × age | 1 | 65.00 | 0.39 | .532 | .555 | .01 | [.00, 1.00] |
| R Ventral Striatum | **modality** | **1** | **65.00** | **8.91** | **.004** | **.014*** | **.12** | **[.02, 1.00]** |
| | age | 1 | 65.00 | 0.33 | .569 | .569 | .01 | [.00, 1.00] |
| | modality × age | 1 | 65.00 | 0.90 | .346 | .378 | .01 | [.00, 1.00] |

*Note*. Type III Analysis of Variance table with Satterthwaite's method. Highlighted in grey are the significant effects after correction for multiple comparisons using the FDR method. L = left, R = right, ROI = region of interest, RPE = reward prediction error, NumDF = numerator degrees of freedom, DenDF = denominator degrees of freedom, *p* unc. = uncorrected *p*-value, $\eta^2_p$ = partial eta-squared effect sizes, CI = confidence interval, suppl. = supplementary. Signif. codes: 0 '***' 0.001 '**' 0.01 '*' 0.05 '.' 0.1 ' ' 1. Interpretation of effect sizes: $\eta^2_P < .01$ very small, $.01 \leq \eta^2_P < .06$ small, $.06 \leq \eta^2_P < .14$ medium, $\eta^2_P \geq .14$ large (Field, 2013).

#### B.8 ROI analyses for value processing mask

**Figure B8**

*Depiction of value network used for ROI analyses*

**Value network**

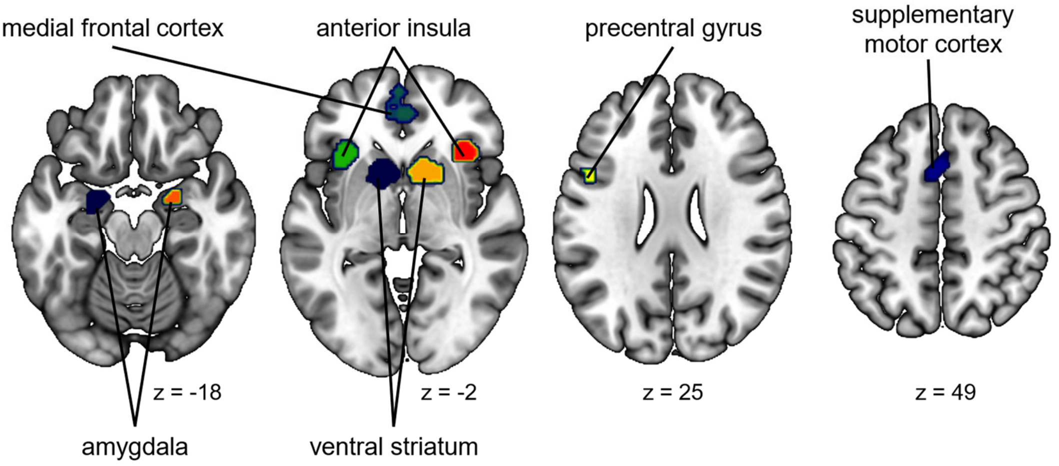


*Note.* The figure shows the clusters included in the value network mask. ROI = region of interest.

We did not observe any significant effects of modality, age, or the interaction between modality and age in the value network ROI.
